# BAZ1A, an Imitation Switch (ISWI) protein, interacts and facilitates the recruitment of E2F1 to activate the E2F transcription program

**DOI:** 10.1101/2024.11.20.624462

**Authors:** Bidisha Prajapati, Abhishek Chowdhury, Kaval Reddy Prasasvi, Lakshay Garg, Kumaravel Somasundaram

## Abstract

ISWI (*I*mitation *Swi*tch) family of chromatin remodeling complexes mobilize nucleosomes to regulate DNA-template associated functions. Analysis of glioblastoma (GBM) transcriptome datasets revealed that BAZ1A (**<u>B</u>romodomain <u>A</u>djacent to <u>Z</u>inc Finger Domain <u>1A</u>**) is a highly expressed ISWI family member. RNAi-based depletion or inhibition by a small molecule inhibited the survival/migration of glioma cells and glioma stem-like cells (GSCs), sensitized glioma cells to Temozolomide but did not affect normal astrocytes. BAZ1A-silencing arrested cells in the G1 phase and induced apoptosis. While GO analysis of BAZ1A regulated transcriptome showed enrichment of “cell cycle” related terms, GSEA showed a depletion of the “HALLMARK_E2F_TARGETS” gene set with the highest significance, indicating a nonfunctional E2F transcription program. BAZ1A silencing and overexpression experiments confirmed the requirement of BAZ1A for the E2F transcription program by identifying E2F1 as the primary target. ChIP experiments revealed that BAZ1A and E2F1 bind to the E2F1 promoter. ChIP-ReChIP demonstrated that BAZ1A-bound chromatin fragments are enriched for E2F1 protein on the E2F1 promoter at specific sites. SMARCA1/5, ATPase subunits of the ISWI family, also exhibited binding to the E2F1 promoter. BAZ1A depletion decreased the DNaseI sensitivity of E2F1 binding regions of the E2F1 promoter. Co-immunoprecipitation experiments revealed an interaction between BAZ1A and E2F1 and the presence of an E2F1-BAZ1A-SMARCA1/5 complex. Finally, BAZ1A silencing inhibited glioma tumor growth in an orthotopic xenograft mouse model. Our results demonstrate that BAZ1A containing ISWI complex recruits E2F1 to activate the E2F transcription program, thus promoting G1-S progression and highlighting BAZ1A as a potential therapeutic target.

## Introduction

Chromatin remodeling is the dynamic alteration of the structure and organization of the chromatin, which plays a crucial role in regulating gene expression and controlling access to the DNA by various cellular machinery. Chromatin remodelers utilize energy from ATP hydrolysis to move, reposition, or eject nucleosomes, thereby altering the accessibility of DNA to transcription factors, RNA polymerases, and other regulatory proteins. Thus, by modulating the organization of chromatin, they can activate or silence specific genes, influencing crucial biological processes such as cellular differentiation, DNA replication, DNA repair, and genomic stability [1].

Chromatin remodeler proteins are broadly classified into several families based on their structure, composition, and mechanism of action, which include SWI/SNF (Switch/Sucrose Non-Fermentable), ISWI (Imitation SWI), CHD (Chromodomain Helicase DNA-binding) and INO80 (INOsitol-requiring mutant 80) [2]. Deregulation or mutations in chromatin remodeler proteins have been implicated in various human diseases, including cancer. Understanding the activity and regulation of chromatin remodeler proteins is thus critical for deciphering the molecular mechanisms underlying normal cellular functions and disease development. Though other families of chromatin remodelers have received considerable attention, evolutionary conserved ISWI group complexes have received the same far less, both mechanistically and functionally. Typically, ISWI complexes facilitate the development of premature nucleosomes into mature octameric nucleosomes, maintain nucleosome spacing at comparatively constant distances, and are crucial in transcriptional regulation ([3], [4], [5], and [6]) DNA damage response and repair, DNA recombination, and many other aspects of cell physiology ([7], [8] and [9]).

Glioblastoma (GBM) is the most common adult brain tumor with a poor prognosis ([10]). The role of SWI/SNF in GBM and other brain tumor pathogenesis is well established ([11]). BRG1, the catalytic subunit of the SWI/SNF complex, downregulates TXNIP, anegative regulator of glycolysis, to maintain a stem-like state of glioma-initiating cells ([12]). SMARCD3, a SWI/SNF complex subunit, has been shown to regulate DAB1-Reelin to promote medulloblastoma metastasis ([13]). However, the role of ISWI complex in GBM pathogenesis has yet to be studied. In this study, we demonstrate that BAZ1A is the most upregulated ISWI complex protein in GBM. We show that BAZ1A promotes glioma cell proliferation and survival by facilitating the E2F transcription program. We also found that BAZ1A maintains *in vivo* glioma tumor growth, thus making it a potential therapeutic target.

## Materials and Methods

### Cell lines and culturing conditions

The GBM cell lines included in this study are - LN229 (a kind gift from late Dr. Abhijit Guha,University of Toronto, Canada), U87 (Sigma Aldrich (U.S.A)), and U-87 MG-Luc2 that express luciferase gene luc2 (gifted by Dr. Vooijs (Maastricht University Medical Center, Netherlands)). NHA-hTERT-E6/E7 was obtained from Dr. A. Guha’s laboratory. HEK293T (obtained from ECACC) and all other cell lines were grown in Dulbecco’s modified Eagle’s medium (DMEM) supplemented with 10% Fetal Bovine Serum (FBS), penicillin (Sigma, U.S.A), streptomycin (Sigma, U.S.A) and gentamicin (SRL). The cells were maintained in the humidified incubator at 37°C and 5% CO2. Primary tumor Glioma stem-like cells (GSCs), MGG8 was kindly provided by Dr. Wakimoto (Massachusetts General Hospital, Boston). SK-1035 was a generous gift from Dr. Santosh Kesari (University of California, San Diego). GSCs were cultured as neurospheres in ultra-low attachment plates (Corning, U.S.A) in Neurobasal medium (Invitrogen, U.S.A.) supplemented with three mmol/L L-Glutamine (Invitrogen, U.S.A.), basic fibroblast growth factor (bFGF; 20 ng/ml, Promega), epidermal growth factor (EGF; 20 ng/ml; Promega), 1X B27 supplement (Invitrogen, U.S.A.), 0.5X N-2 (Invitrogen, U.S.A.), 2 μg/mL Heparin (Sigma, U.S.A.), penicillin (Sigma, U.S.A.), gentamicin, and streptomycin (Sigma, U.S.A.). Neurospheres were dissociated into single cells and passaged every seven days using a chemical dissociation kit (Catalog# 05707, STEMCELL technologies, U.S.A.). Fresh neurobasal medium (containing all the supplements) was added every alternative day. RT-qPCR was utilized to test for mycoplasma contamination. The absence of mycoplasma in all cell lines is confirmed using the EZdetect PCR Kit for Mycoplasma Detection from HiMedia.

### Other materials and methods

The details regarding the following methods are given in the Supplementary information. Differential expression analysis, Correlation analysis, RNA sequencing, Gene set enrichment analysis, Plasmids and siRNAs, Lentivirus preparation and infection, Sphere formation assay, Limiting dilution assay, RNA isolation, cDNA synthesis, and RT-qPCR, Proliferation assay using MTT, Viability assay, Chemosensitivity assay, Colony formation assay, Migration assays, Cell cycle analysis by flow cytometry, BrdU incorporation assay using flow cytometry, Annexin V-FITC (Fluorescein isothiocyanate) and PI staining, Luciferase assay, Immunoblotting, Immunoprecipitation assays, Chromatin Immunoprecipitation (ChIP), ChIP ReChIP or sequential Chromatin Immunoprecipitation, DNaseI hypersensitivity assay, Orthotopic xenograft mouse model, *In vivo* imaging, Cryo-sectioning of fixed mouse brain, Immunohistochemistry (IHC), Hematoxylin and Eosin staining (H and E).

## Results

### BAZ1A is highly expressed in GBM, and its expression is under the control of the Wnt/β-catenin signaling pathway

To explore the importance of ISWI chromatin remodeler complex in GBM, we investigated the transcript level expression of all regulatory (BAZ1A, BAZ1B, BAZ2A, BAZ2B, BPTF, CECR2, CHRAC1, POLE3, RBBP4, RBBP7, and RSF1) and catalytic (SMARCA1 and SMARCA5) subunits of ISWI group in GBM transcriptome datasets (n=10). Our results showed that BAZ1A (**<u>B</u>romodomain <u>A</u>djacent To <u>Z</u>inc Finger Domain <u>1A</u>)** is the most highly upregulated gene across the majority of the datasets analyzed (**Figure 1A, B; Supplementary Tables 1 and 2**). Hence, we chose to study the role of BAZ1A as a critical regulator of the ISWI family in GBM. We also found that the BAZ1A transcript level is significantly higher in low-grade gliomas (LGG) in addition to GBM compared to control brain samples (**Figure 1C; Supplementary Figure 1A**). The protein level of BAZ1A is highly elevated in GBM, as seen by immunohistochemical analysis (**Figure 1D and E**) and mass spectrometry analysis (**Figure 1F**). Pan-cancer data analysis revealed that BAZ1A is highly expressed in most cancers at transcript and protein levels (**Supplementary Figure 1B and C**).

**Figure 1:**
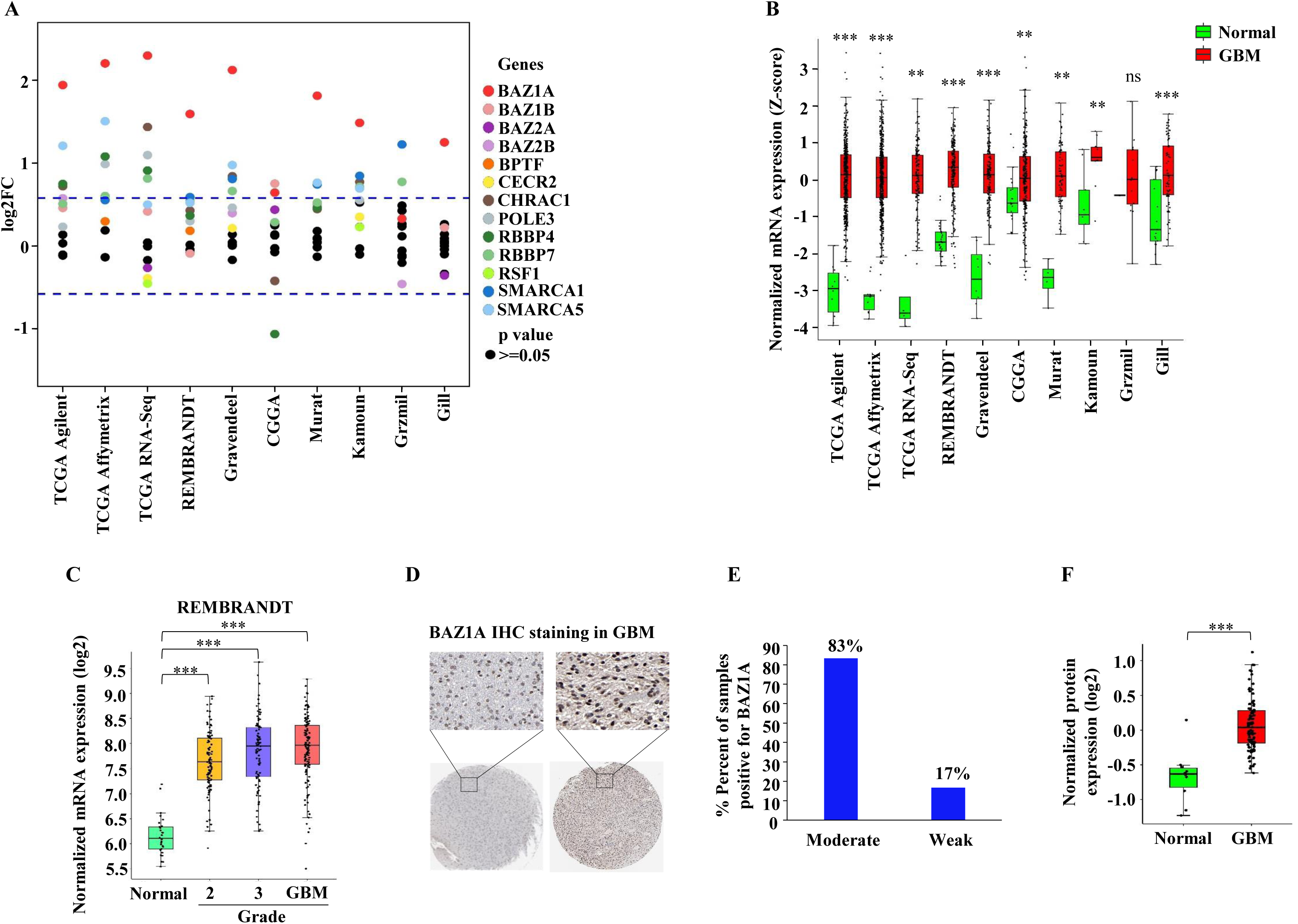
BAZ1A is a highly upregulated ISWI group chromatin remodeler protein in GBM. **A**. The transcript level expression of 13 ISWI group proteins was analyzed in ten different GBM datasets (**GlioVis**: http://gliovis.bioinfo.cnio.es/ and **CGGA**: http://www.cgga.org.cn/.). The log2 fold change (GBM-Normal) values were plotted for each gene. **B**. Transcript level expression of BAZ1A in ten different GBM datasets compared to control (**GlioVis**: http://gliovis.bioinfo.cnio.es/ and **CGGA**: http://www.cgga.org.cn/). **C**. Grade-wise transcript expression of BAZ1A in different grades of GBM and control samples in REMBRANDT dataset (**GlioVis**: http://gliovis.bioinfo.cnio.es/). **D** and **E**. BAZ1A protein expression (IHC staining) in glioma patient tissue samples and their quantification (Data obtained from The Human Protein Atlas, https://www.proteinatlas.org/). **F**. BAZ1A protein expression in control and GBM samples (Data obtained from CPTAC, https://proteomics.cancer.gov/programs/cptac). Student’s t-test was performed, where *:p<0.05,**:p<0.01,***:p<0.001, ns: non-significant.

In order to dissect the mechanism behind BAZ1A’s higher expression in glioma, we investigated the impact of treatment of glioma cells with pharmacological inhibitors ofdifferent oncogenic signaling pathways on BAZ1A expression. We found that treatment with FH535, a Wnt/β-Catenin pathway inhibitor, significantly decreased the BAZ1A transcript andprotein levels (**Figure 2A and B**). BAZ1A transcript and protein levels showed a concentration-dependent and time-dependent reduction in FH535-treated LN229 cells (**Figure 2C, D, E, and F**) and U87 glioma cells (**Figure 2G, H, I, and J**). Further, treatment with Lithium chloride (LiCl), an activator of the Wnt pathway, showed a substantial increase in the BAZ1A transcript and protein levels in LN229 and U87 cells (**Figure 2K, L, M, and N**). Knockdown of β-catenin, a key member of the Wnt pathway, showed a significant decrease in BAZ1A transcript and protein levels in LN229 and U87 cells (**Figure 2O, P, Q, and R**). Silencing LEF1, the heterodimeric partner of β-Catenin, also reduced BAZ1A transcript and protein levels in LN229 and U87 cells (**Figure 2S, T, U, and V**). These results collectively confirm that the Wnt/β-catenin signaling pathway is responsible for the higher expression of BAZ1A in glioma. Based on these results, we conclude that BAZ1A, a component of the ISWI complex, is highly expressed in GBM, and its expression is regulated by the Wnt/β-catenin signaling pathway.

**Figure 2:**
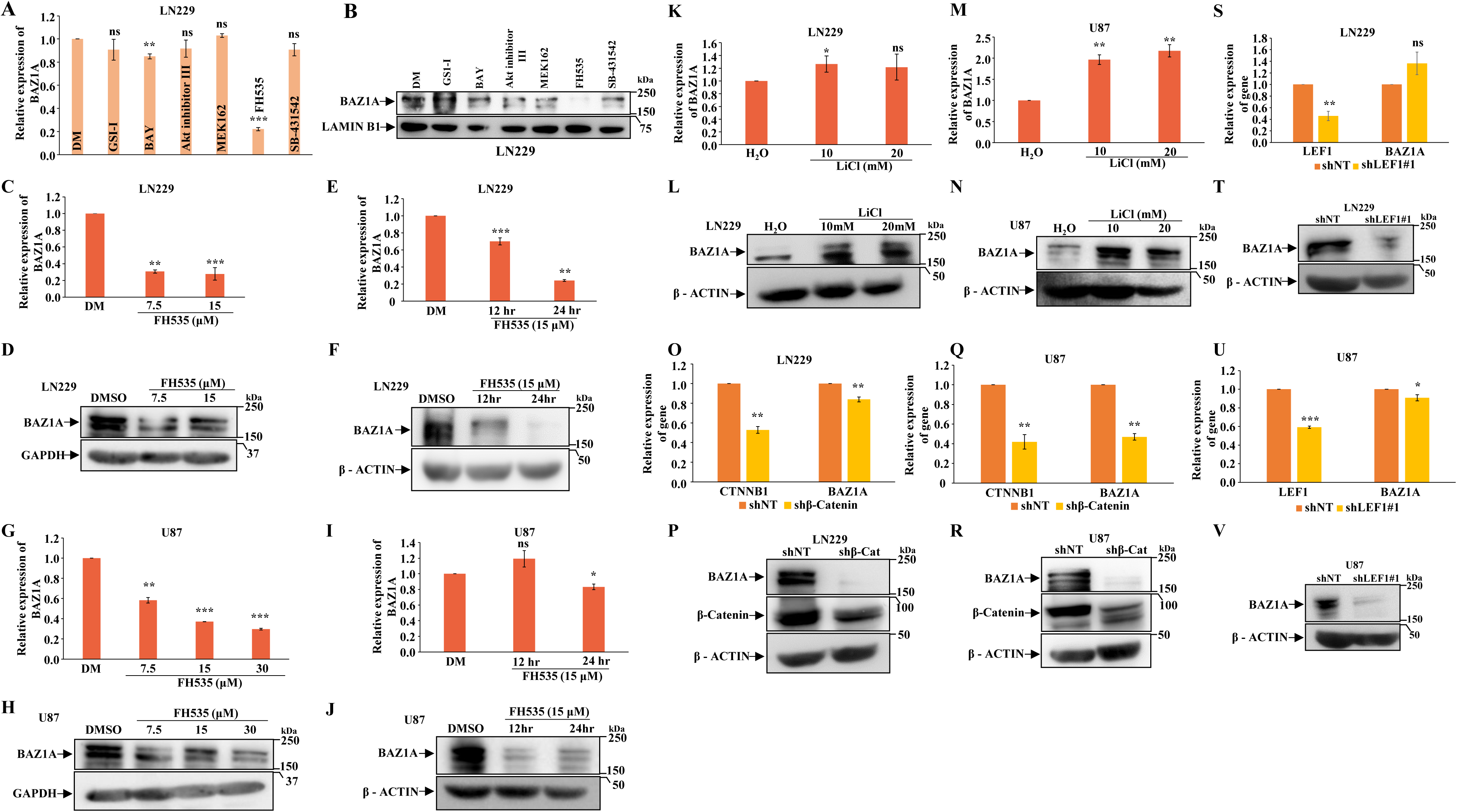
Wnt/β-catenin signaling pathway regulates the expression of BAZ1A in GBM. **A** and **B**. BAZ1A transcript (**A**) and protein level (**B**) expression was measured after 24 hours of inhibitor (six different inhibitors) treatment of LN229 cells. **C** and **D**. RT-qPCR and western blot of BAZ1A after 24 hours of FH535 treatment of LN229 cells. **E** and **F**. RT-qPCR and western blot of BAZ1A after 12 and 24 hours of FH535 treatment of LN229 cells. **G** and **H**. RT-qPCR and western blot of BAZ1A after 24 hours of FH535 treatment of U87 cells. **I** and **J**. RT-qPCR and western blot of BAZ1A after 12 and 24 hours of FH535 treatment of U87 cells. **K** and **L**. RT-qPCR and western blot of BAZ1A after 24 hours of LiCl treatment of LN229 cells. **M** and **N**. RT-qPCR and western blot of BAZ1A after 24 hours of LiCl treatment of U87 cells. **O** and **P**. RT-qPCR and western blot of BAZ1A and β-catenin after silencing of β-catenin in LN229 cells. **Q** and **R**. RT-qPCR and western blot of BAZ1A and β-catenin after silencing of β-catenin in U87 cells. **S** and **T**. RT-qPCR and western blot of BAZ1A after silencing of LEF1 in LN229 cells. **U** and **V**. RT-qPCR and western blot of BAZ1A after silencing of LEF1 in U87 cells. Student’s t-test was performed, where *:p<0.05,**:p<0.01,***:p<0.001, ns: non-significant.

### BAZ1A promotes cell survival, and its inhibition sensitizes glioma cells to chemotherapy

Next, we investigated the functional role of highly expressed BAZ1A in glioma. BAZ1A expression was inhibited using two individual shRNAs, and the impact on various cancer cell-related phenotypes was examined. BAZ1A silencing in LN229 and U87 glioma cells reduced cell proliferation, colony formation, and migration (**Figure 3A, B, C, and D; Supplementary Figure 2A, B, C, and D)**. BAZ1A silenced glioma cells showed G1 arrest with a significant reduction in the S phase (**Figure 3E; Supplementary Figure 2E)** and elevated apoptosis (**Figure 3F; Supplementary Figure 2F).** Glioma stem-like cells (GSCs) that exist in small proportions initiate and maintain the tumor and contribute to resistance to therapy and recurrence in GBM ([19] and [20]). BAZ1A silencing inhibited the growth of patient-derived GSCs, as seen by neurosphere growth assay and limiting dilution assay (**Figure 3G and H; Supplementary Figure 2G and H).** In contrast, BAZ1A silencing failed to inhibit the growth of normal human astrocytes **(Figure 3I and J**).

**Figure 3:**
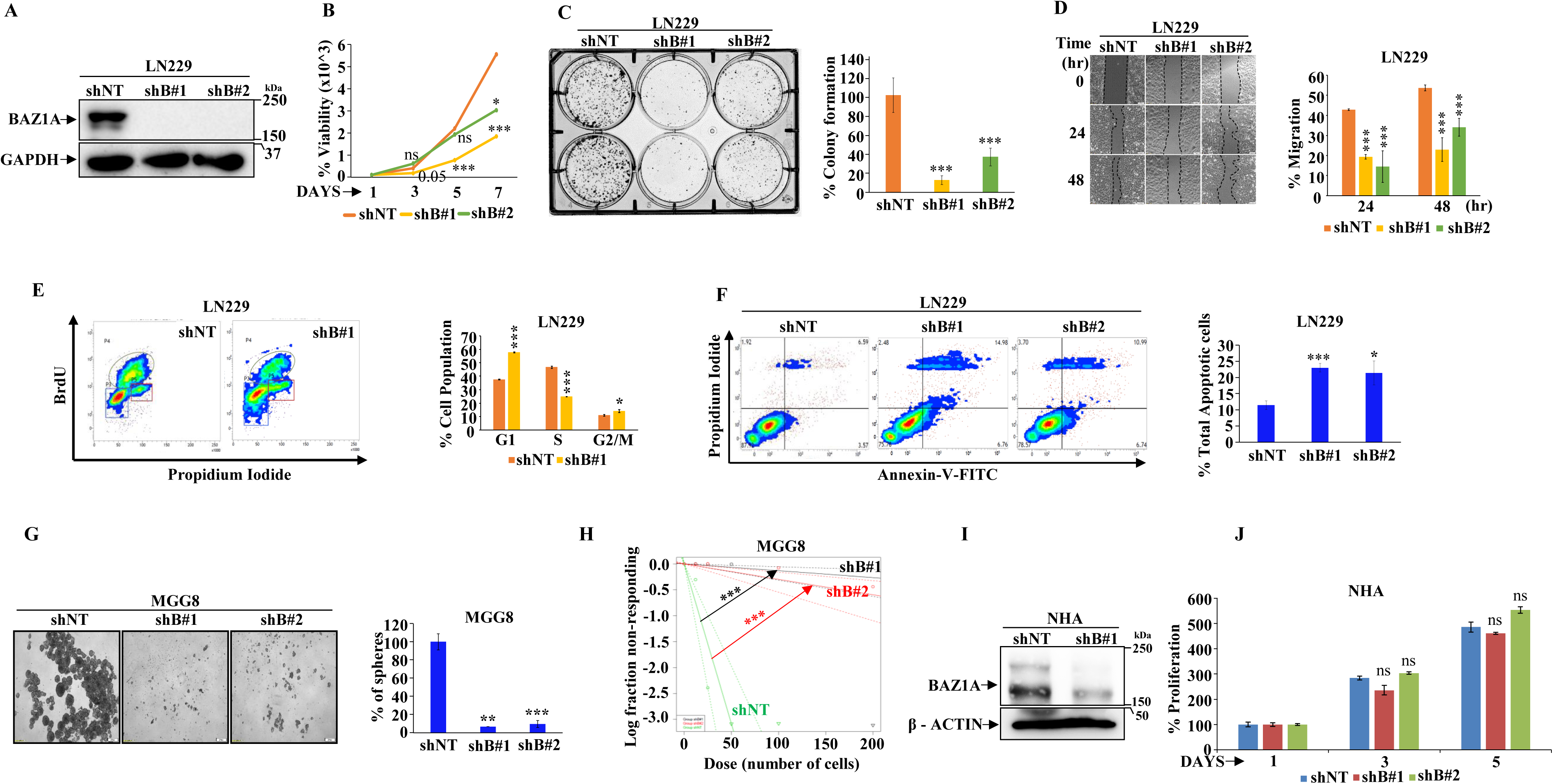
BAZ1A promotes cell survival, cell cycle, migration and inhibits apoptosis. **A**. Immunoblot depicting shRNA mediated silencing of BAZ1A with two individual shRNAs (shB#1 and shB#2) in LN229 cells. **B**. Cell viability assay with BAZ1A-depleted and control LN229 cells. **C**. Colony formation assay after silencing of BAZ1A in LN229 cells and their quantification. **D**. Scratch assay or migration assay after knockdown of BAZ1A in LN229 cells and their quantification. **E**. Cell cycle assay using BrdU and propidium iodide after silencing of BAZ1A in LN229 cells and their quantification. **F**. Annexin-V-FITC assay of BAZ1A-depleted LN229 cells and their quantitation. **G**. Neurosphere growth assay in MGG8 after depletion of BAZ1A and their quantification. **H**. Limiting dilution assay of MGG8 cells after knockdown of BAZ1A. χ2 test was performed for pair-wise differences. **I**. Immunoblot of BAZ1A silencing in NHA cells. **J**. Proliferation assay using MTT after silencing of BAZ1A in NHA. Student’s t-test was performed, where *:p<0.05,**:p<0.01,***:p<0.001, ns: non-significant.

Since Temozolomide (TMZ) chemotherapy is prescribed for GBM, we tested the sensitivity of BAZ1A silenced cells to TMZ. BAZ1A silencing sensitized the glioma cells to TMZ as seen by reduced proliferation, deregulated cell cycle, and induction of massive apoptosis in LN229 and U87 glioma cells (**Supplementary Figure 3A, B, C, D, E, and F).** BAZ1A inhibition by Cpd-2, a pharmacological inhibitor ([21]), also inhibited glioma cell growth as seen by reduced proliferation (**Supplementary Figure 4A),** cell cycle arrest in the G1 phase (**Supplementary Figure 4B)**, and induction of apoptosis (**Supplementary Figure 4C**) and sensitized glioma cells to temozolomide chemotherapy (**Supplementary Figure 4D).** Thus, we conclude that BAZ1A is essential for glioma cell survival, and its absence or inhibition sensitizes glioma cells to temozolomide chemotherapy.

### BAZ1A is essential for the activation of the E2F transcriptional program

Towards deciphering the mechanistic insight into BAZ1A functions, diferentially expressed genes (**Supplementary Table 3**) between GBM samples with higher (BAZ1A^high^) and lower (BAZ1A^low^) BAZ1A transcript levels were subjected to GO and GSEA analysis. GO analysis showed an enrichment of “Cell_cycle,” and other related terms. Similarly, GSEA also demonstrated the significant enrichment of cell cycle-related gene sets such as “HALLMARK_G2M_CHECKPOINT”, “HALLMARK_E2F_TARGETS”, and “HALLMARK_MITOTIC_SPINDLE” (**Supplementary Figure 5A, B and C**) indicating a possible role for BAZ1A in the regulation of the cell cycle.

Next, total RNA-Seq of BAZ1A silenced cells was obtained to earthen the BAZ1A regulated transcriptome to confirm the above results and identify the BAZ1A targets. BAZ1A silenced cells showed 2899 differentially expressed genes (DEGs; downregulated -1533 and upregulated -1366) (**Figure 4A; Supplementary Table 4**). GO analysis of DEGs showed enrichment of terms - cell cycle, cell division, and chromosome (**Figure 4B; Supplementary Table 5**). GSEA showed a significant depletion of several cancer-related gene sets, with “HALLMARKS_E2F_TARGETS” showing the highest significance (**Figure 4C, D; Supplementary Figure 6A; Supplementary Table 6**), suggesting the presence of a defective E2F transcriptional program in BAZ1A silenced cells. Based on these observations, we hypothesized that BAZ1A may be required for E2F-dependent transcription. To test this possibility, we measured the transcript levels of E2F target genes in glioma cells with BAZ1A silencing or exogenous overexpression. While BAZ1A silencing significantly decreased the transcript levels of E2F target genes in LN229 (**Figure 4E)** and U87 glioma cells (**Supplementary Figure 5D),** BAZ1A overexpressing glioma cells showed enhanced transcript levels of E2F target genes (**Figure 4F; Supplementary Figure 5E).** Next, we measured the transcription from E2F1-dependent promoter-reporters in cells with altered BAZ1A levels. In BAZ1A silenced LN229 and U87 cells, luciferase activity from CDK4 promoter-Luc and cMYC promoter-Luc is significantly reduced (**Figure 4G and H; Supplementary Figure 6B and C, respectively**). Since the E2F1 promoter is activated by E2F family transcription factors, we also measured the impact of BAZ1A silencing on the transcription of the E2F1 promoter-Luc reporter. The transcription from E2F1 promoter-Luc is also reduced significantly in BAZ1A-silenced LN229 and U87 cells (**Figure 4I; Supplementary Figure 6D, respectively**). In contrast, exogenous overexpression of BAZ1A increased the activity of all three reporters in both cells (**Figure 4J, K, and L; Supplementary Figure 6E, F, and G, respectively**). From these results, we conclude that BAZ1A is crucial for activating the E2F transcriptional program in glioma.

**Figure 4:**
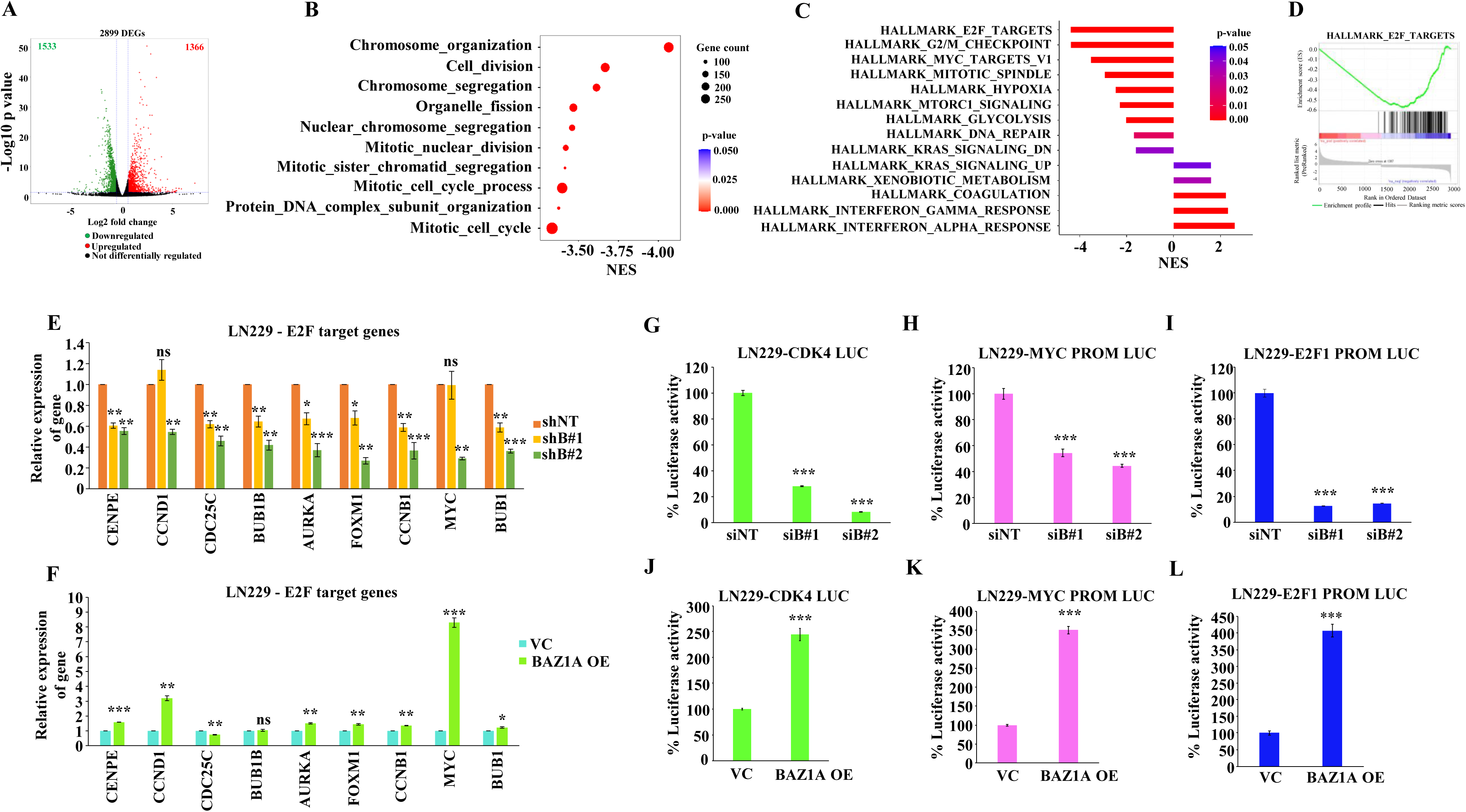
BAZ1A regulates E2F transcriptional program. **A**. Volcano plot showing differentially regulated genes in BAZ1A silenced LN229 cells. **B**. Gene ontology (GO) analysis of DEGs in BAZ1A silenced LN229 cells. **C**. Gene set enrichment analysis (GSEA) analysis of DEGs in BAZ1A-silenced LN229 cells. **D**. Enrichment plot showing “HALLMARK_E2F_TARGTES” gene set in BAZ1A silenced LN229 cells. **E**. RT-qPCR of E2F target genes in BAZ1A depleted LN229 cells. **F**. RT-qPCR of E2F target genes in BAZ1A overexpressed condition in LN229 cells. Cells were harvested after 24 hours of transfection of BAZ1A overexpression construct. **G**, **H** and **I**. CDK4, MYC and E2F1 promoter luciferase activity measurement in BAZ1A-silenced LN229 cells. Cells were harvested after 72 hours of transfection of siRNA and luciferase constructs. **J**, **K** and **L**. CDK4, MYC and E2F1 promoter luciferase activity measurement in BAZ1A overexpressed LN229 cells. Cells were harvested after 24 hours of transfection of BAZ1A overexpression and luciferase constructs. Student’s t-test was performed, where *:p<0.05,**:p<0.01,***:p<0.001, ns: non-significant.

### BAZ1A interaction with E2F1 recruits E2F1 to E2F1 promoter to initiate E2F transcription program

Next, we sought to identify the key target of BAZ1A in controlling the E2F transcription program. A family of eight E2F genes encoding transcription factors controls the E2F transcription program to orchestrate the cell cycle progression through the G1-S-G2/M phases ([22]). E2F family consists of three activators (E2F1, E2F2, and E2F3), repressors (E2F4, E2F5, and E2F6), and atypical repressors (E2F7, and E2F8). Both activators and repressors bind to the same enhancer sequences but result in transcriptional activation and repression, respectively ([22] and [23]). Our results thus far show that the E2F transcription program is defective in BAZ1A-silenced cells.

To identify BAZ1A’s target, we examined the expression status of E2F genes in BAZ1A-silenced cells. GSEA output showed that many E2F target gene transcripts are repressed in BAZ1A-silenced cells (**Figure 5A**). A closer look at the repressed genes revealed that four E2F genes - E2F1, E2F2, E2F7, and E2F8 - were downregulated at the transcript level (**Figure 5B and C**), while all others were not regulated. The transcript downregulation of E2F1, E2F2, E2F7, and E2F8 was independently validated in BAZ1A silenced glioma cells (LN229 and U87) and GSC lines (SK-1035 and MGG8) (**Figure 5D; Supplementary Figure 7A, B and C**). Transcript level correlation revealed a significant positive correlation between BAZ1A and all four E2Fs in GBM patient data (**Figure 5E)**, which was further validated in publicly available GBM datasets (n=24) (**Supplementary Table 7 and 8)**, suggesting that BAZ1A may influence the expression of these four E2F genes positively. Since the activator type of E2F transcription factors initiates the E2F transcription program, we focused on E2F1 and E2F2. The protein level expression of the above four E2Fs in BAZ1A-silenced conditions showed a consistent downregulation of the E2F1 protein in both LN229 and U87 cell lines (**Figure 5F and G**) indicating that E2F1 is likely a critical target of BAZ1A. Further, the exogenous overexpression of BAZ1A in glioma cells also increased the E2F1 transcript (**Figure 5H and I**) and E2F1 protein (**Figure 5J and K**) levels. BAZ1A silencing decreased the luciferase activity activated specifically by exogenously expressed E2F1 (**Figure 5L**). Taken together, these results indicate that E2F1, the bonafide activating type of E2F, could be the main target of BAZ1A in regulating the E2F transcription program.

**Figure 5:**
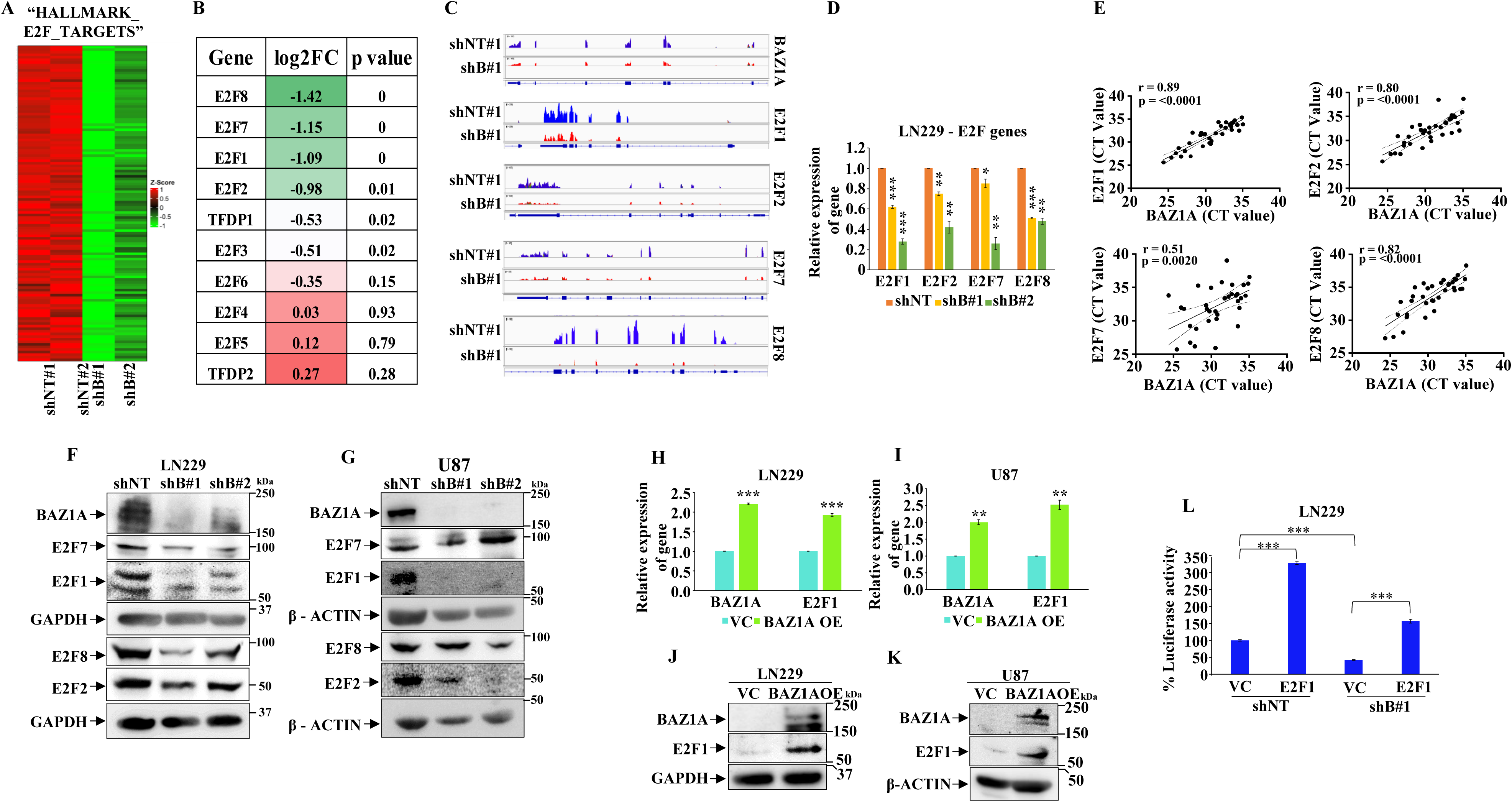
E2F1 gene expression is regulated by BAZ1A. **A**. Heatmap of E2F target genes from the “HALLMARK_E2F_TARGETS” gene set after BAZ1A knockdown in LN229 cells. **B**. Table showing log2fold change and p values of a list of E2F genes and TFDP1 and TFDP2 genes after BAZ1A silencing in LN229 cells. **C**. Genome tracks showing the peaks of read counts of BAZ1A, E2F1, E2F2, E2F7, and E2F8 in BAZ1A-silenced LN229 cells. Integrative Genomics Viewer (IGV) is used. **D**. RT-qPCR analysis of E2F1, E2F2, E2F7, and E2F8 transcripts in BAZ1A-depleted LN229 cells. **E**. Transcript level correlation of BAZ1A with E2F1, E2F2, E2F7, and E2F8 in our in-house GBM patient samples. CT values obtained from RT-qPCR are plotted. **F** and **G**. Immunoblot analysis of E2F1, E2F2, E2F7 and E2F8 protein in BAZ1A silenced LN229 and U87 cells. **H** and **I**. RT-qPCR analysis of E2F1 transcript in BAZ1A overexpressed condition in LN229 and U87 cells. Cells were harvested after 24 hours of transfection of BAZ1A overexpression construct. **J** and **K**. Immunoblot analysis of E2F1 protein in BAZ1A overexpressed condition in LN229 and U87 cells. Cells were harvested after 24 hours of transfection of BAZ1A overexpression construct. **L**. E2F1 promoter luciferase activity measurement in control and BAZ1A-silenced cells after exogenous E2F1 overexpression. Student’s t-test was performed, where *:p<0.05,**:p<0.01,***:p<0.001, ns: non-significant.

BAZ1A binds to acetylated lysine residues through its bromodomain and regulates the ATPase activity of ACF complex by binding to SMARCA1/5, thus remodeling the chromatin to regulate transcription and DNA damage repair ([24] and [21]). Among E2F1, E2F2, and E2F3, the activating type of E2Fs, E2F1 is mainly required for initial entry into the cell cycle as it binds to its own promoter and that of E2F2 and E2F3 to initiate the E2F transcription program ([25]). Hence, we hypothesized that BAZ1A may facilitate E2F1 binding to its own promoter. First, we investigated BAZ1A’s binding to the E2F1 promoter. ChIP with BAZ1A antibody showed that BAZ1A protein occupies the E2F1 promoter region with varying enrichment levels at multiple sites (**Figure 6A and B)**. We also found that BAZ1A binds to multiple sites on the promoters of E2F2 and E2F3 (**Supplementary Figures 8A, B, C, and D**). ChIP with E2F1 antibody showed enrichment of E2F1 protein on E2F1 promoter at specific sites (C, D, and G) located near predicted E2F1 binding sites (**Figure 6A and 6C**). Since the sites C, D, and G on the E2F1 promoter were bound by BAZ1A and E2F1, we focussed on these sites for further investigations. ISWI and SWI/SNF complexes selectively mediate the binding of specific transcription factors by enhancing chromatin accessibility ([6]). To test the possibility of BAZ1A binding to E2F1 promoter at specific regulatory sites (C, D, and G) that may enhance chromatin accessibility for E2F1 binding, we performed a quantitative DNaseI sensitivity assay. This revealed that the sites C, D, and G of E2F1 are less sensitive to DNaseI in BAZ1A silenced cells (**Figure 6D, E, and F**), suggesting that BAZ1A bound chromatin is more accessible for E2F1 protein.

**Figure 6:**
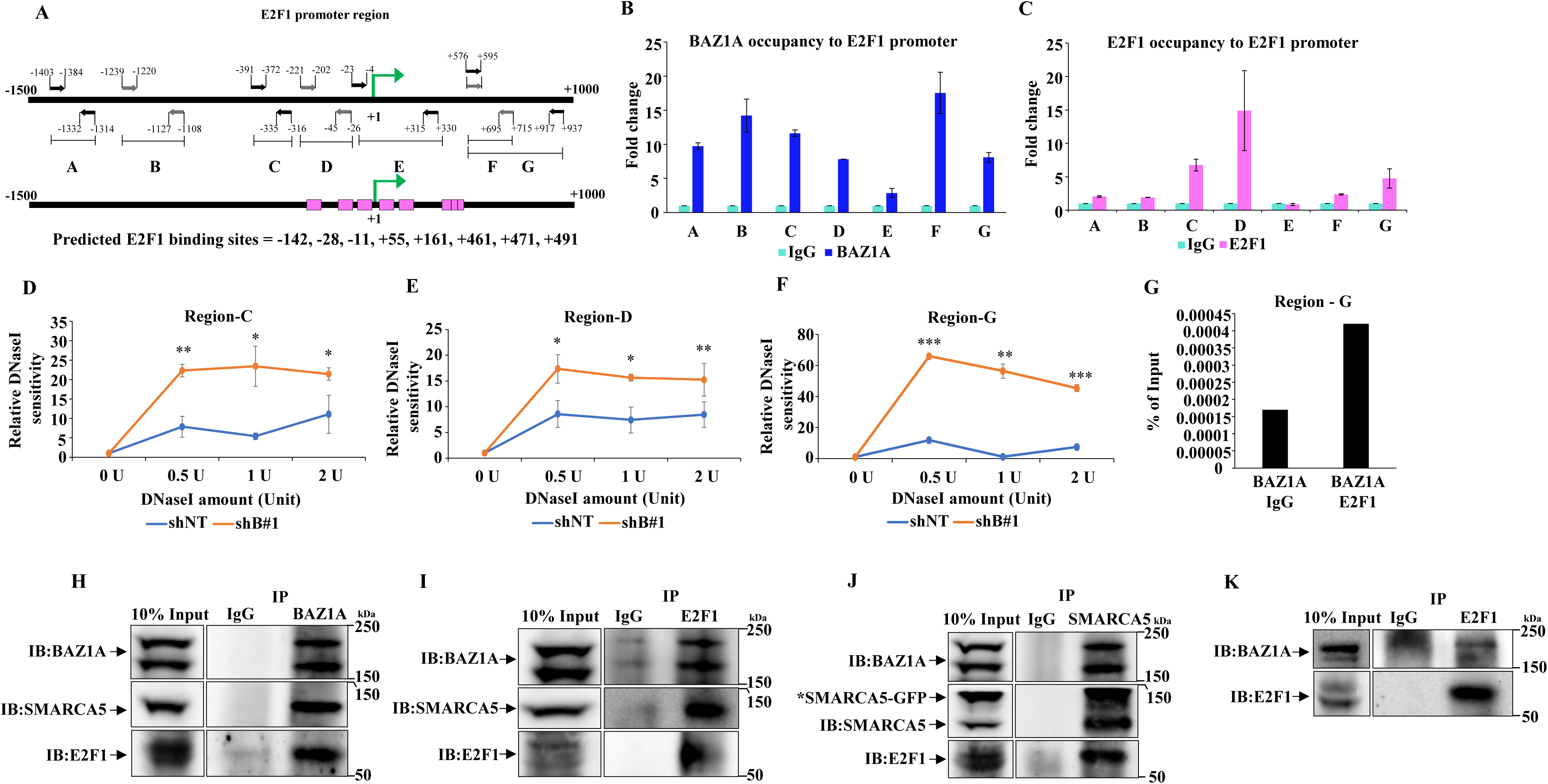
BAZ1A occupies E2F1 promoter region, remodels it, and help recruit E2F1. **A**. Schematic of E2F1 promoter region and predicted E2F1 binding sites on it. **B**. ChIP followed by qPCR with BAZ1A antibody to E2F1 promoter region. IgG was used as a control. **C**. ChIP followed by qPCR with E2F1 antibody to E2F1 promoter region. IgG was used as a control. **D, E and F**. DNaseI hypersensitivity assay of BAZ1A-silenced LN229 cells with an increasing amount of DNaseI. qPCR analysis of E2F1 promoter region C, D, and G. **G**. ChIP with BAZ1A followed by ReChIP with E2F1 or IgG. ChIP ReChIP assay followed by PCR of region G of E2F1 promoter. **H** and **I**. HEK293T cells were transfected with BAZ1A-eGFP and HA-E2F1 and harvested after 24 hours for whole cell lysate preparation. Immunoprecipitation for BAZ1A (**H**) or E2F1 (**I**) was performed with the lysates with specific antibodies, which were then resolved by SDS-PAGE, and immunoblotting was performed with indicated antibodies. 10% of inputs are indicated. **J**. HEK293T cells were transfected with BAZ1A-eGFP, HA-E2F1, and SMARCA5-eGFP and harvested after 24 hours for whole cell lysate preparation. Immunoprecipitation for SMARCA5 was performed on the lysates with SMARCA5 antibody, which were then resolved by SDS-PAGE, and immunoblotting was performed with indicated antibodies. 10% of inputs are indicated. **K**. LN229 cells were serum starved for 48 hours, released in 12% serum-containing medium, and harvested at the G1-S phase transition point. Lysates were prepared for immunoprecipitation with E2F1 antibody, which was then resolved by SDS-PAGE, and immunoblotting was performed with indicated antibodies. 10% of inputs are indicated. Student’s t-test was performed where *:p<0.05,**:p<0.01,***:p<0.001, ns: non-significant.

BPTF, a component of the NURF ISWI complex, interacts and recruits c-MYC to its target genes ([26]). Hence, we decided to investigate the possibility of a coordinated binding by BAZ1A and E2F1 proteins to the E2F1 promoter, which may facilitate the E2F1 gene transcription. Hence, we performed ChIP-ReChIP or sequential ChIP. As shown before, BAZ1A ChIP showed efficient binding to sites C, D, and G of the E2F1 promoter (**Supplementary Figure 8E, F, G**). ReChIP of BAZ1A bound chromatin fragments with E2F1 antibody showed that BAZ1 bound E2F1 promoter region is enriched for E2F1 at the site G of the E2F1 promoter (**Figure 6G**) but not at other sites (**data not shown**). The above results indicate a co-occupancy of BAZ1A and E2F1 on the E2F1 promoter at a specific site. This result raises the possibility of a potential interaction between BAZ1A and E2F1 proteins and a possible recruitment of E2F1 by chromatin-bound BAZ1A. We performed co-immunoprecipitation in the HEK293T cell line upon overexpressing BAZ1A (BAZ1A-eGFP) and E2F1 (HA-E2F1). BAZ1A immunoprecipitation showed an enrichment of E2F1 and SMARCA5 proteins (**Figure 6H**). Similarly, E2F1 immunoprecipitation showed an enrichment of BAZ1A and SMARCA5 proteins (**Figure 6I)**. In addition, SMARCA5 or SMARCA1 immunoprecipitations also brought down BAZ1A and E2F1 in the same immune complex (**Figure 6J; Supplementary Figure 8H**). In LN229 glioma cells that are entering into the S phase (16 hrs after the release from G1 arrest by serum starvation), E2F1 immunoprecipitation showed an enrichment of BAZ1A, thus confirming the interaction between endogenous E2F1 and BAZ1A proteins under endogenous conditions (**Figure 6K**). We also confirmed that the E2F1 promoter is enriched for SMARCA1 and SMARCA5 proteins at sites C, D, and G (**Supplementary Figure 8I and J**). Based on these results, we conclude that E2F1, BAZ1A, and SMARCA1/5 exist in a complex, and chromatin-bound BAZ1A recruits E2F1 onto the E2F1 promoter. Collectively, we propose that a trimeric complex containing BAZ1A, SMARCA1/5, and E2F1 proteins on the E2F1 promoter initiates the E2F1 gene transcription.

### BAZ1A is essential for glioma growth *in vivo*

To test the importance of BAZ1A on glioma tumor growth, we tested the ability of U87 cells silenced for BAZ1A in an intracranial mouse model. U87-Luc/shBAZ1A cells formed significantly smaller tumors than U87-Luc/shNT cells (**Figure 7A, B, and C**). The mice with U87-Luc/shBAZ1A tumors showed increased survival (**Figure 7D**). Further, U87-Luc/shBAZ1A tumors showed reduced expression of E2F1 compared to U87-Luc/shNT tumors (**Figure 7E**). In addition, the staining of Ki67, a proliferation marker, is significantly lesser in U87-Luc/shBAZ1A tumors than in U87-Luc/shNT tumors (**Figure 7F**). These results confirm that BAZ1A is essential for glioma tumor growth.

**Figure 7:**
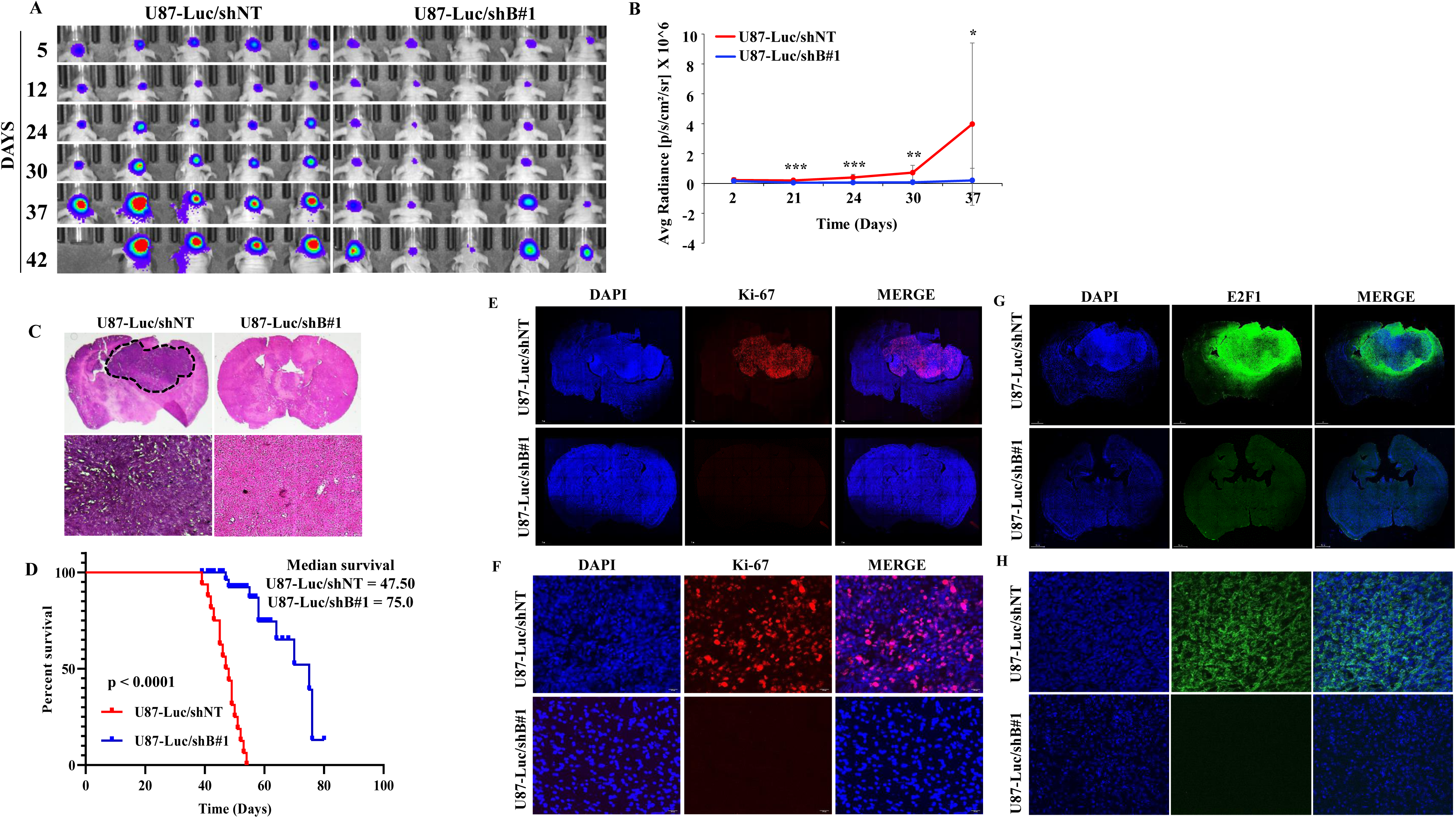
BAZ1A is essential for glioma tumor growth *in vivo*. **A** and **B**. Representative images of *in vivo* tumor bioluminescence of the U87-Luc/shNT and U87-Luc/shB#1 mice and the measurement of the bioluminescence is plotted. **C**. Hematoxylin and Eosin staining of the tumor-bearing U87-Luc/shNT and U87-Luc/shB#1 mice brain sections (0.7X and 40X). **D**. Kaplan–Meier survival graph reflecting the survival of U87-Luc/shNT and U87-Luc/shB#1 tumor-bearing mice (n=16). Mantel-Cox log-rank test was performed for the significance. **E** and **F**. Immunofluorescence images of the whole brain (**E**, 10X) and the tumor sections (**F**, 40X) of the brain of U87-Luc/shNT and U87-Luc/shB#1 mice stained for proliferation marker Ki-67 and DAPI. **G** and **H**. Immunofluorescence images of the whole brain (**G**, 10X) and the tumor sections (**H**, 40X) of the brain of U87-Luc/shNT and U87-Luc/shB#1 mice stained for E2F1 and DAPI. Student’s t-test was performed, where *:p<0.05,**:p<0.01,***:p<0.001, ns: non-significant.

## Discussion

Altered chromatin remodeling plays an important role in cancers. ISWI chromatin remodeling complexes are essential for cell survival and have been shown to play a critical role in tumorigenesis. While the role played by ISWI family members through genetic and epigenetic changes in many cancers is well documented, its role in glioma development is scanty ([27]). We found that BAZ1A, one of the regulatory subunits of the ISWI family, is highly expressed in GBM. We show that BAZ1A containing ISWI complex controls glioma growth by remodeling the chromatin at the E2F1 promoter, facilitating E2F1 binding to activate the E2F transcription program.

While BPTF and RSF1, members of the ISWI complex, have been able to regulate glioma through different signaling pathways, the precise nature of chromatic remodeling during gliomagenesis and the mechanism behind their overexpression is not established ([28] and [29]). Our comprehensive study demonstrates that BAZ1A is the most upregulated ISWI complex member in GBM in several glioma datasets. Interestingly, the higher expression of BAZ1A is seen in low-grade gliomas, suggesting a potential role for BAZ1A in the initial glioma transformation event. This is further strengthened by our finding that BAZ1A promotes cell cycle progression by facilitating the E2F transcription program through chromatin remodeling (see later for details). We also found that the oncogenic Wnt/β-Catenin signaling pathway controls the higher expression of BAZ1A. BAZ1A promoted glioma cell proliferation and survival as either shRNA-mediated downregulation of its expression or inhibition by its bromodomain-specific inhibitor inhibited growth and induced programmed cell death.

Investigation of differentially expressed genes in BAZ1A silenced cells aptly identified an enrichment of cell cycle-related terms indicating the crucial role of BAZ1A in regulating glioma cell proliferation and survival. Deeper studies revealed a non-functional E2F transcription program in BAZ1A silenced or inhibited cells that supports G1 phase arrest, as seen in BAZ1A silenced cells. A well-orchestrated regulation by the E2F family of transcription factors controls the cell cycle progression through the G1-S phase of the cell cycle ([22]). The activator E2Fs (E2F1, E2F2 and E2F3) play a major role in executing E2F transcription program as they activate S phase-specific genes during the G1-S phase transition and the atypical repressors (E2F7 and E2F8) during late S phase which in turn shut off the expression of activator E2Fs and E2F targets to shut of E2F transcription program thus allowing cells to progress to G2 phase. Our study found that E2F1, the bonafide activator type of E2F, is the direct target of BAZ1A. While BAZ1A silencing inhibited E2F1 transcript levels, its overexpression induced E2F1 expression. Mechanistically, we found that BAZ1A facilitates E2F1 expression by multiple ways: 1) BAZ1A binds to the E2F1 promoter at multiple sites and remodels the chromatin as seen by the increased sensitivity of E2F1 promoter to DNAase1; 2) ChIP experiments revealed binding of BAZ1A and E2F1 proteins at specific sites on E2F1 promoter; 3) Co-immunoprecipitation experiments revealed an interaction between BAZ1A and E2F1 in an E2F1-BAZ1A-SMARCA1/5 protein complex; and 4) Chromatin boundBAZ1A recruited E2F1 to a specific site on the E2F1 promoter. Thus, our study demonstratesthe fundamental role of BAZ1A in regulating the cell cycle by facilitating E2F1 recruitment to promote the E2F transcription program. We propose a model (**Figure 8**) wherein Wnt/β-catenin controlled BAZ1A regulates the E2F transcription program in regulating the cell cycle progression.

**Figure 8:**
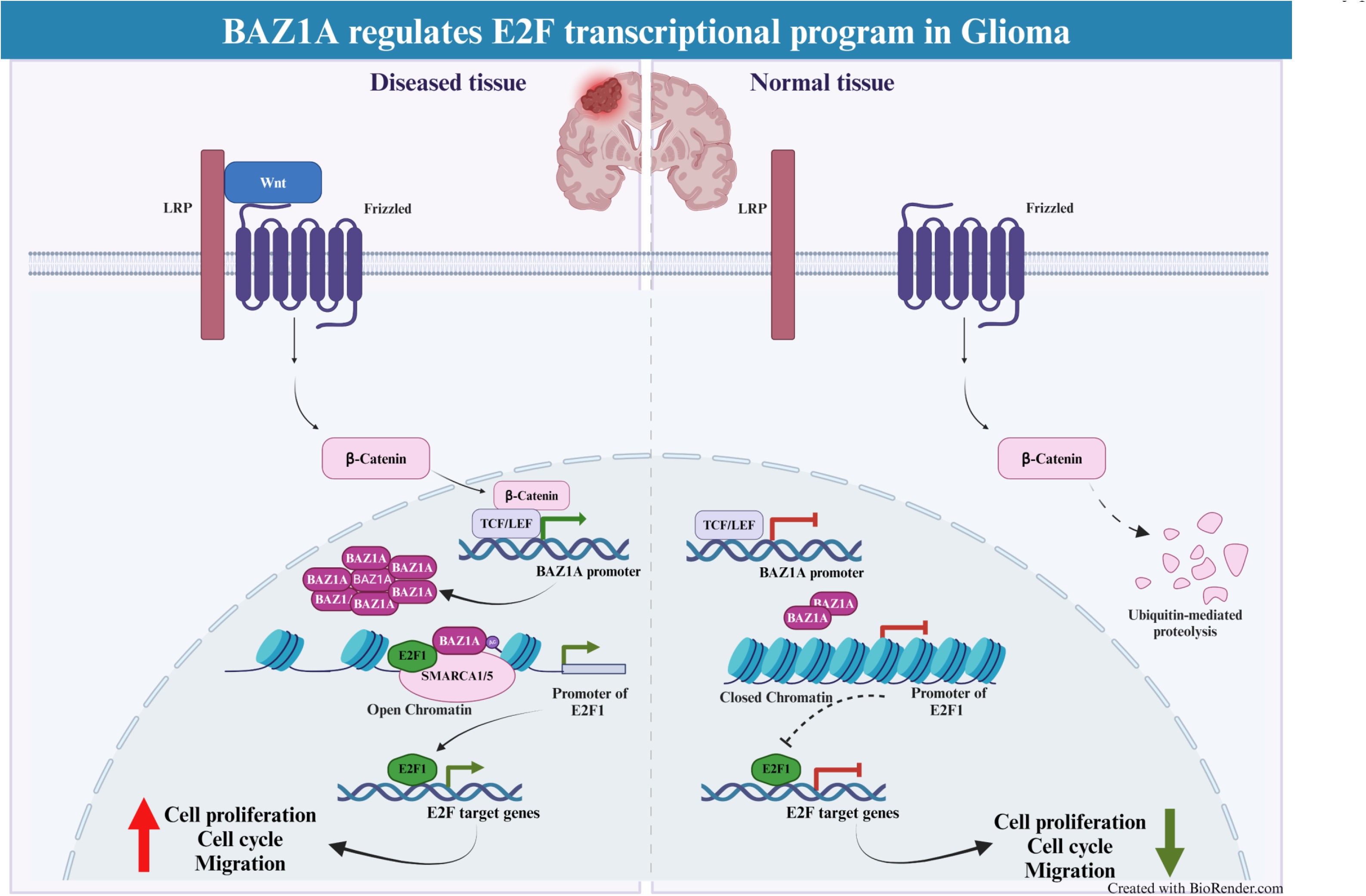
Schematic representation of molecular mechanism of BAZ1A in glioma.

Bromodomain-containing proteins are recognized as potential targets for cancer. Specific inhibitors targeting the bromodomain of BAZ1A have been developed ([21]). We found BAZ1A silencing inhibited glioma tumors in an intracranial mouse model, supporting the critical role of BAZ1A in the E2F transcription program and cell cycle progression. Collectively, our study establishes the crucial role of BAZ1A containing ISWI remodeling complex in glioma and proposes BAZ1A as a therapeutic target for GBM.

## Acknowledgments

The results published here are, in whole or part, based upon data generated by The Cancer Genome Atlas pilot project established by the NCI and NHGRI. Information about TCGA andthe investigators and institutions that constitute the TCGA research network can be found at http://cancergenome.nih.gov/. We acknowledge the shRNA consortium (Dr. Subba Rao), IISc, India, for shRNA constructs. BP acknowledges the IISc fellowship, and KRP acknowledges DBT for fellowship. KS acknowledges CEFIPRA, DBT, DST, and CSIR (Govt. of India) for research grants. Infrastructure supported by DST FIST, DBT-IISc partnership program, and UGC is acknowledged.

## Author contributions

BP planned and carried out all the experiments; AC performed all bioinformatic analyses; KRPand BP carried out animal experiments; LG carried out experiments related to the cell cycle; KS prepared the manuscript, planned the study, and led the project.

## Abbreviations

E2F1: E2F transcription factor 1
BAZ1A: Bromodomain Adjacent to Zinc Finger Domain 1A
ISWI: Imitation Switch
SMARCA: SWI/SNF related, matrix associated, actin dependent regulator of chromatin, Subfamily A
GBM: Glioblastoma

## Notes

### Competing Interest Statement

The authors have declared no competing interest.

