## Supplementary Figures for "BAZ1A, an Imitation Switch (ISWI) protein, interacts and facilitates the recruitment of E2F1 to activate the E2F transcription program"

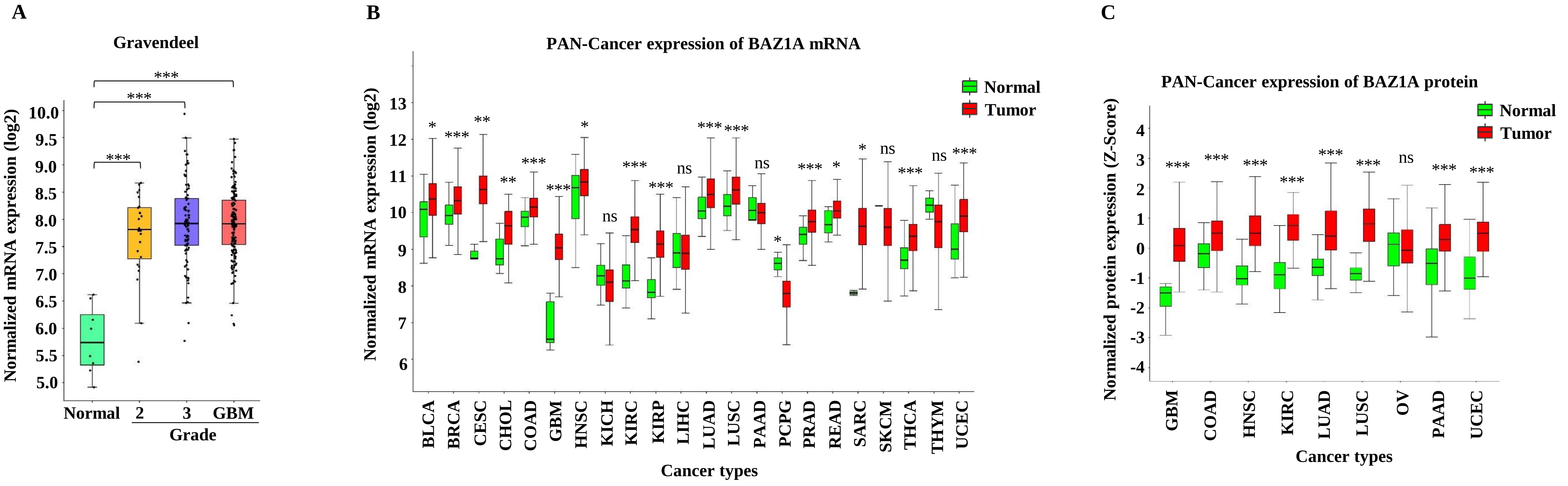
A
Gravendeel
Normalized mRNA expression (log2)
10.0
9.5
9.0
8.5
8.0
7.5
7.0
6.5
6.0
5.5
5.0
***
***
***
Normal
2
3
GBM
Grade
B
PAN-Cancer expression of BAZ1A mRNA
Normalized mRNA expression (log2)
13
12
11
10
9
8
7
6
Normal
Tumor
*
***
**
**
***
***
*
ns
***
***
ns
***
***
ns
*
***
*
*
ns
***
ns
***
BLCA
BRCA
CESC
CHOL
COAD
GBM
HNSC
KICH
KIRC
KIRP
LIHC
LUAD
LUSC
PAAD
PCPG
PRAD
READ
SARC
SKCM
THCA
THYM
UCEC
Cancer types
C
PAN-Cancer expression of BAZ1A protein
Normalized protein expression (Z-Score)
4
3
2
1
0
-1
-2
-3
-4
Normal
Tumor
***
***
***
***
***
***
ns
***
***
GBM
COAD
HNSC
KIRC
LUAD
LUSC
OV
PAAD
UCEC
Cancer types

**Supplementary Figure 1**: **BAZ1A is a highly upregulated ISWI group chromatin remodeler protein in GBM. A**. Grade-wise transcript expression of BAZ1A in different grades of GBM and control samples in Gravendeel dataset (**GlioVis**: http://gliovis.bioinfo.cnio.es/). **B**. Transcript level expression of BAZ1A in pan-cancer data (**TCGA Expression Browser**: https://tools.altiusinstitute.org/tcga/ (Pan-cancer data). Wilcoxon rank-sum test was performed for the significance calculation. **C**. Pan-cancer protein data (CPTAC, https://proteomics.cancer.gov/programs/cptac) showing expression of BAZ1A in different cancer types compared to control samples. Student's t-test was performed, where *:$p<0.05$,**:$p<0.01$,***:$p<0.001$, ns: non-significant.

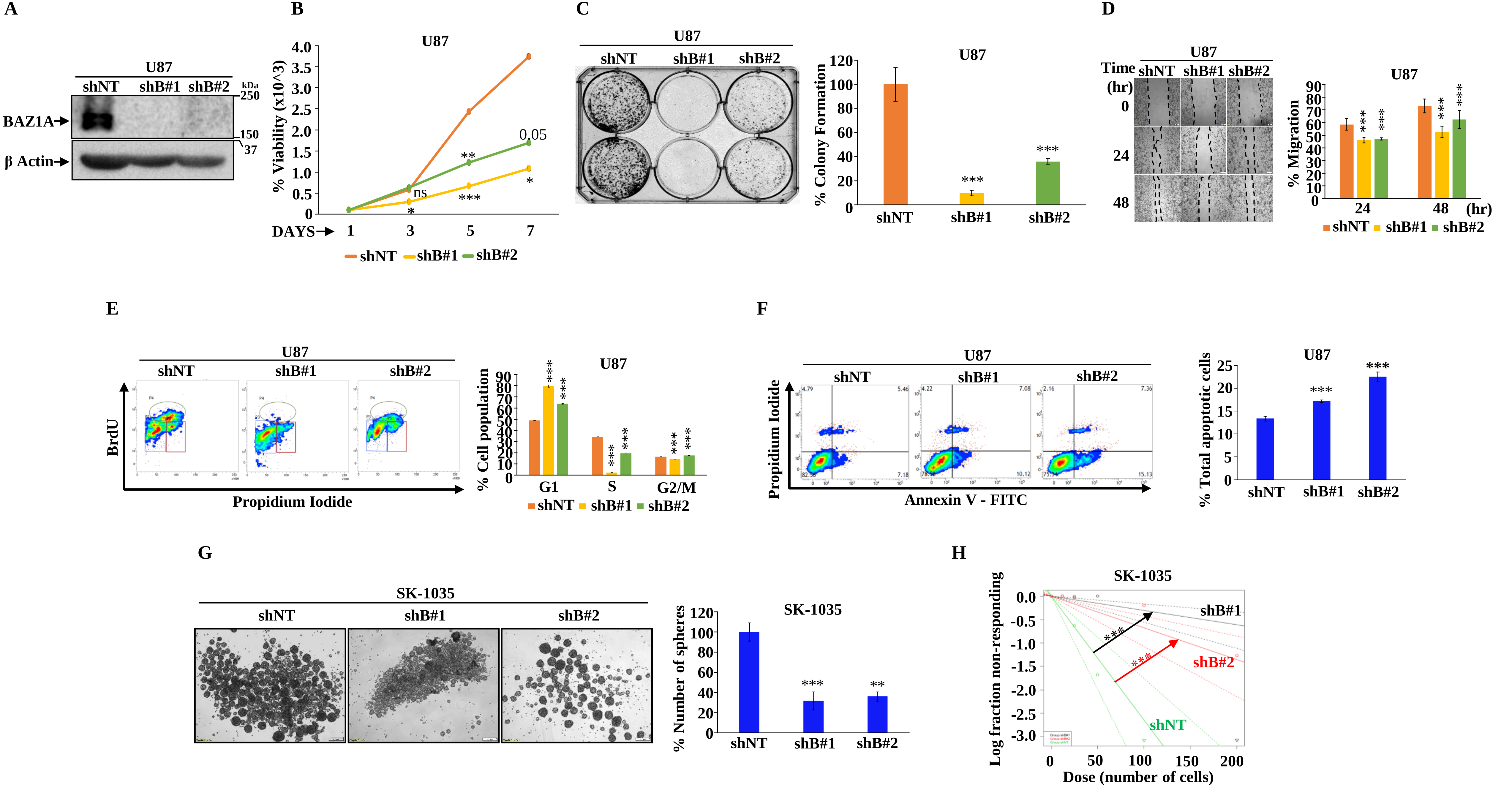
A
U87
shNT shB#1 shB#2
kDa
250
150
37
BAZ1A
β Actin
B
U87
% Viability (x10^3)
DAYS
1 3 5 7
ns
0.05
shNT shB#1 shB#2
C
U87
shNT shB#1 shB#2
% Colony Formation
D
U87
Time (hr)
shNT shB#1 shB#2
0
24
48
% Migration
24 48 (hr)
E
U87
shNT shB#1 shB#2
BrdU
Propidium Iodide
% Cell population
G1 S G2/M
F
U87
shNT shB#1 shB#2
Propidium Iodide
Annexin V - FITC
% Total apoptotic cells
G
SK-1035
shNT shB#1 shB#2
% Number of spheres
H
SK-1035
Log fraction non-responding
Dose (number of cells)
shB#1
shB#2
shNT

**Supplementary figure 2**: **BAZ1A promotes cell survival, cell cycle, migration and inhibits apoptosis.** **A**. Immunoblot depicting shRNA mediated silencing of BAZ1A with two individual shRNAs (shB#1 and shB#2) in U87 cell line. **B**. Cell viability assay with BAZ1A-depleted and control U87 cells. **C**. Colony formation assay after silencing of BAZ1A in U87 cells and their quantification. **D**. Scratch assay or migration assay after knockdown of BAZ1A in U87 cells and their quantification. **E**. Cell cycle assay using BrdU and propidium iodide after silencing of BAZ1A in U87 cells and their quantification. **F**. Annexin-V-FITC assay of BAZ1A depleted U87 cells and their quantitation. **G**. Neurosphere growth assay in SK-1035 cells after depletion of BAZ1A and their quantification. **H**. Limiting dilution assay of SK-1035 cells after knockdown of BAZ1A. $\chi 2$ test was performed for pair-wise differences. Student's t-test was performed, where *:$p<0.05$,**:$p<0.01$,***:$p<0.001$, ns: non-significant.

Supplementary Figure 3; Prajapati et al., 2024

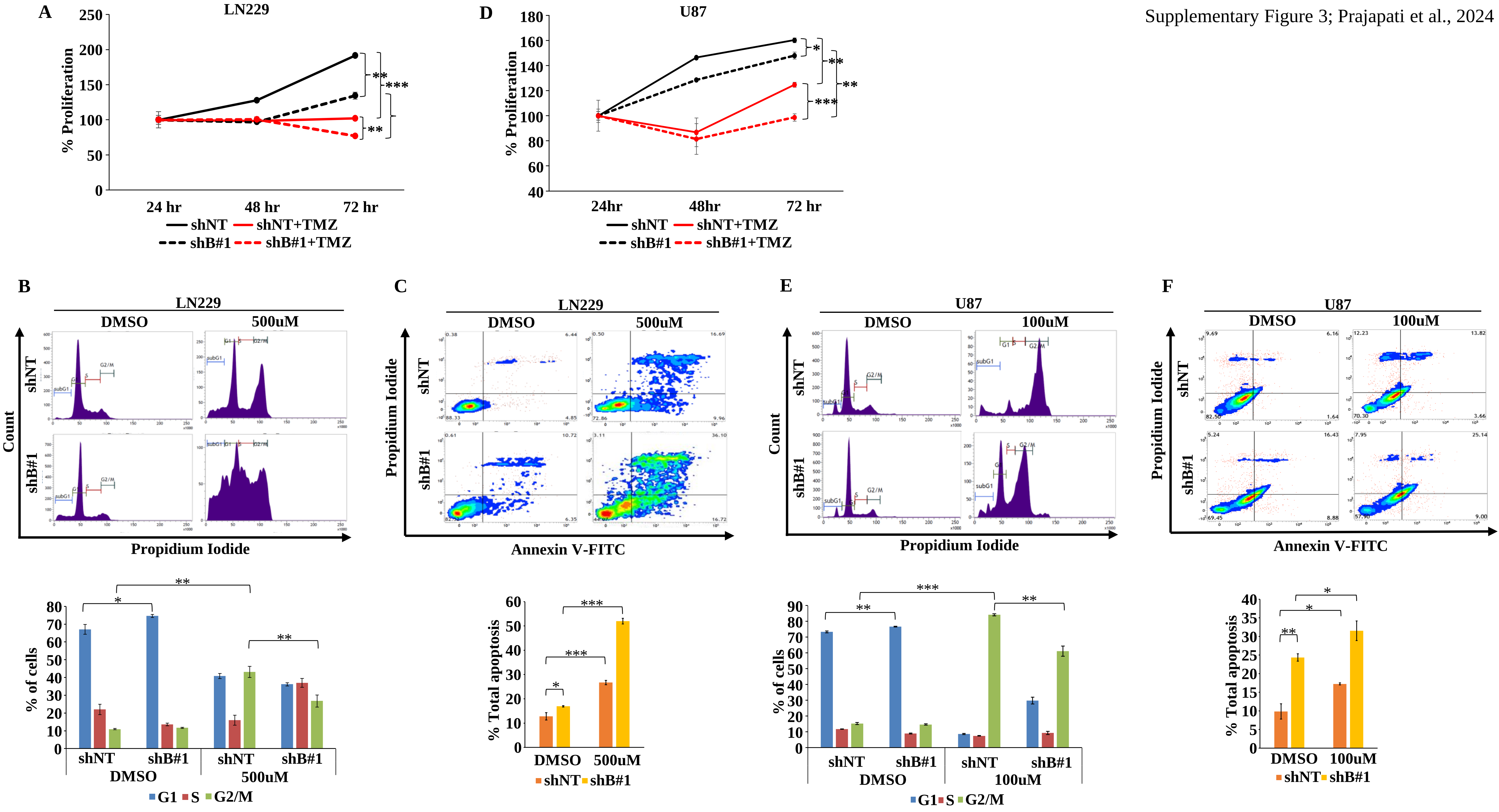

**Supplementary figure 3**: **BAZ1A promotes cell survival, cell cycle, migration and inhibits apoptosis.** **A**. Chemosensitivity assay of BAZ1A compromised LN229 cells. 750 μM of TMZ was used. **B**. BAZ1A-silenced LN229 cells were treated with 500 μM of TMZ and harvested after 72 hours for cell cycle analysis and quantification of cells in G1, S, and G2/M phases has been shown. **C**. BAZ1A-silenced LN229 cells were treated with 500 μM of TMZ and harvested after 72 hours for apoptosis analysis through annexin-V-FITC assay and the percentage of apoptotic cells has been shown. **D**. Chemosensitivity assay of BAZ1A compromised U87 cells. 400 μM of TMZ was used. **E**. BAZ1A-silenced U87 cells were treated with 100 μM of TMZ and harvested after 48 hours for cell cycle analysis and quantification of cells in G1, S, and G2/M phases has been shown. **F**. BAZ1A silenced U87 cells were treated with 100 μM of TMZ and harvested after 48 hours for apoptosis analysis through annexin-V-FITC assay and the percentage of apoptotic cells has been shown. Student's t-test was performed, where *:$p<0.05$,**:$p<0.01$,***:$p<0.001$, ns: non-significant.

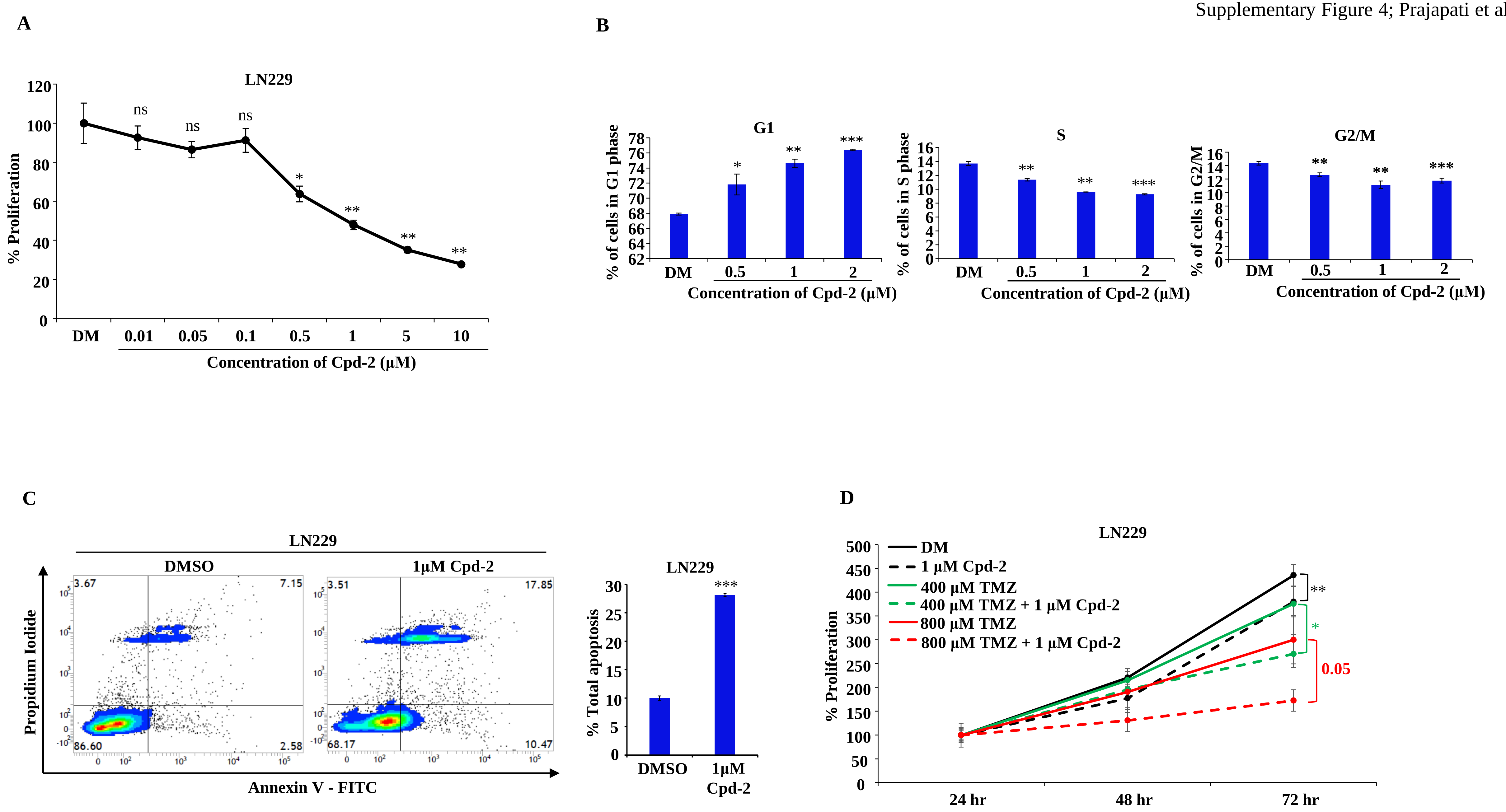
A
LN229
% Proliferation
ns
ns
ns
*
**
**
**
DM 0.01 0.05 0.1 0.5 1 5 10
Concentration of Cpd-2 (μM)
B
G1
% of cells in G1 phase
S
% of cells in S phase
G2/M
% of cells in G2/M
DM 0.5 1 2
Concentration of Cpd-2 (μM)
C
LN229
DMSO
1μM Cpd-2
Propidium Iodide
Annexin V - FITC
3.67
7.15
86.60
2.58
3.51
17.85
68.17
10.47
% Total apoptosis
***
DMSO
1μM Cpd-2
D
LN229
DM
1 μM Cpd-2
400 μM TMZ
400 μM TMZ + 1 μM Cpd-2
800 μM TMZ
800 μM TMZ + 1 μM Cpd-2
% Proliferation
**
*
0.05
24 hr 48 hr 72 hr

**Supplementary figure 4**: **BAZ1A promotes cell survival, cell cycle, migration and inhibits apoptosis. A**. LN229 cell proliferation was measured after 96 hours of treatment with BAZ1A inhibitor Cpd-2. **B**. Cell cycle analysis of LN229 cells after treatment with Cpd-2 for 96 hours. **C**. Apoptosis analysis of LN229 cells via annexin-V-FITC assay upon Cpd-2 inhibitor treatment (1 μM) for 96 hours. **D**. Proliferation assay of LN229 cells after treatment with Cpd-2 (1 μM) and TMZ (400 μM and 800 μM) for different time points. Student’s t-test was performed, where *:$p<0.05$,**:$p<0.01$,***:$p<0.001$, ns: non-significant.

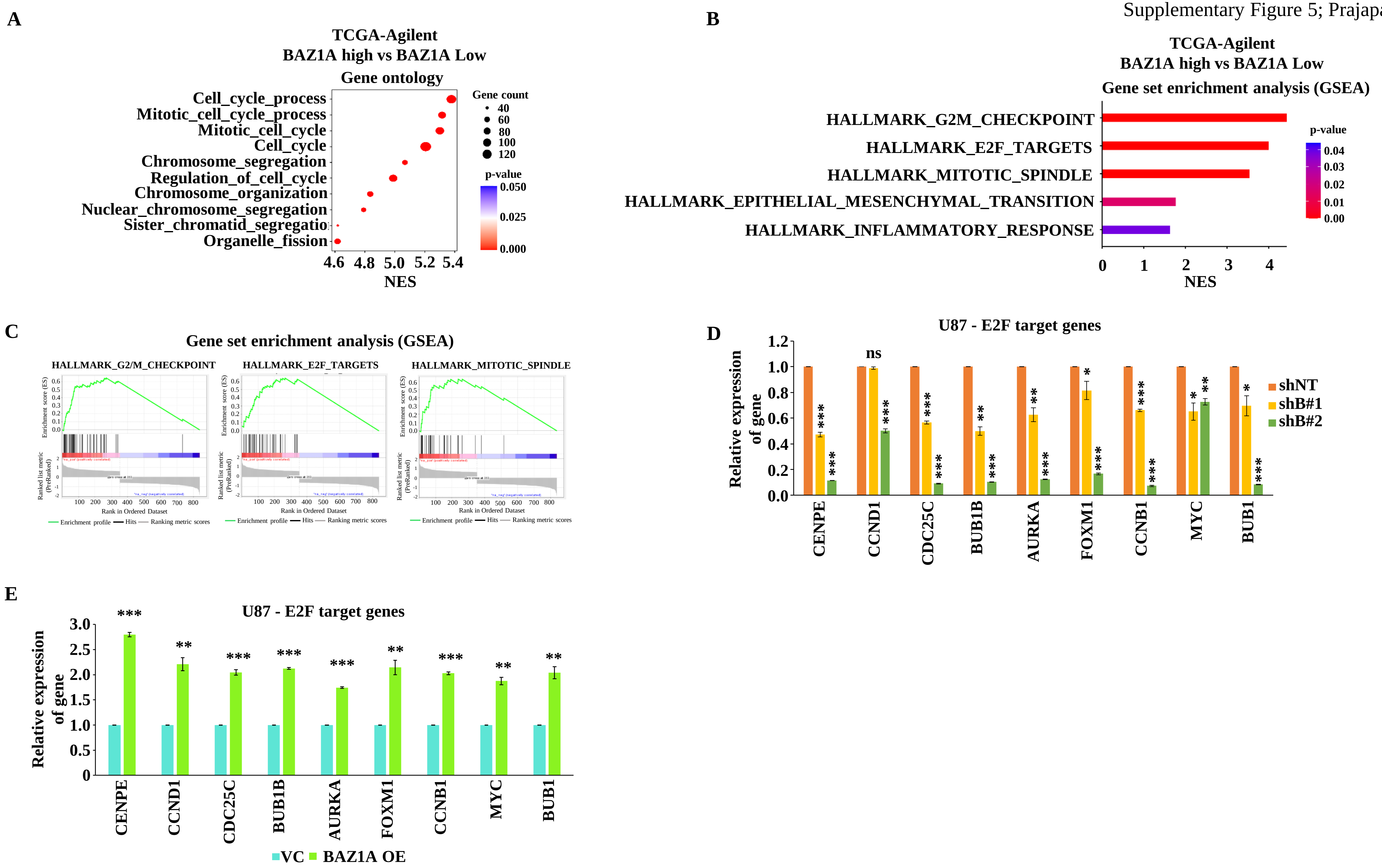
A
TCGA-Agilent
BAZ1A high vs BAZ1A Low
Gene ontology
Cell_cycle_process
Mitotic_cell_cycle_process
Mitotic_cell_cycle
Cell_cycle
Chromosome_segregation
Regulation_of_cell_cycle
Chromosome_organization
Nuclear_chromosome_segregation
Sister_chromatid_segregatio
Organelle_fission
Gene count
40
60
80
100
120
p-value
0.050
0.025
0.000
4.6
4.8
5.0
5.2
5.4
NES
B
TCGA-Agilent
BAZ1A high vs BAZ1A Low
Gene set enrichment analysis (GSEA)
HALLMARK_G2M_CHECKPOINT
HALLMARK_E2F_TARGETS
HALLMARK_MITOTIC_SPINDLE
HALLMARK_EPITHELIAL_MESENCHYMAL_TRANSITION
HALLMARK_INFLAMMATORY_RESPONSE
p-value
0.04
0.03
0.02
0.01
0.00
0
1
2
3
4
NES
C
Gene set enrichment analysis (GSEA)
HALLMARK_G2/M_CHECKPOINT
HALLMARK_E2F_TARGETS
HALLMARK_MITOTIC_SPINDLE
Enrichment score (ES)
Ranked list metric (PreRanked)
Rank in Ordered Dataset
Enrichment profile
Hits
Ranking metric scores
D
U87 - E2F target genes
Relative expression of gene
ns
shNT
shB#1
shB#2
CENPE
CCND1
CDC25C
BUB1B
AURKA
FOXM1
CCNB1
MYC
BUB1
E
U87 - E2F target genes
Relative expression of gene
CENPE
CCND1
CDC25C
BUB1B
AURKA
FOXM1
CCNB1
MYC
BUB1
VC
BAZ1A OE

**Supplementary figure 5**: **BAZ1A regulates E2F transcriptional program.** **A**. Gene ontology (GO) analysis of DEGs obtained from BAZ1A$^{high}$ vs BAZ1A$^{low}$ patient samples. **B**. Gene set enrichment analysis (GSEA) of DEGs obtained from BAZ1A$^{high}$ vs BAZ1A$^{low}$ patient samples. **C**. Plots of the enriched gene sets in GSEA analysis (B). **D**. RT-qPCR of E2F target genes in BAZ1A-silenced condition in U87 cells. **E**. RT-qPCR of E2F target genes in BAZ1A overexpressed condition in U87 cells. Student's t-test was performed, where *:p<0.05,**:p<0.01,***:p<0.001, ns: non-significant.

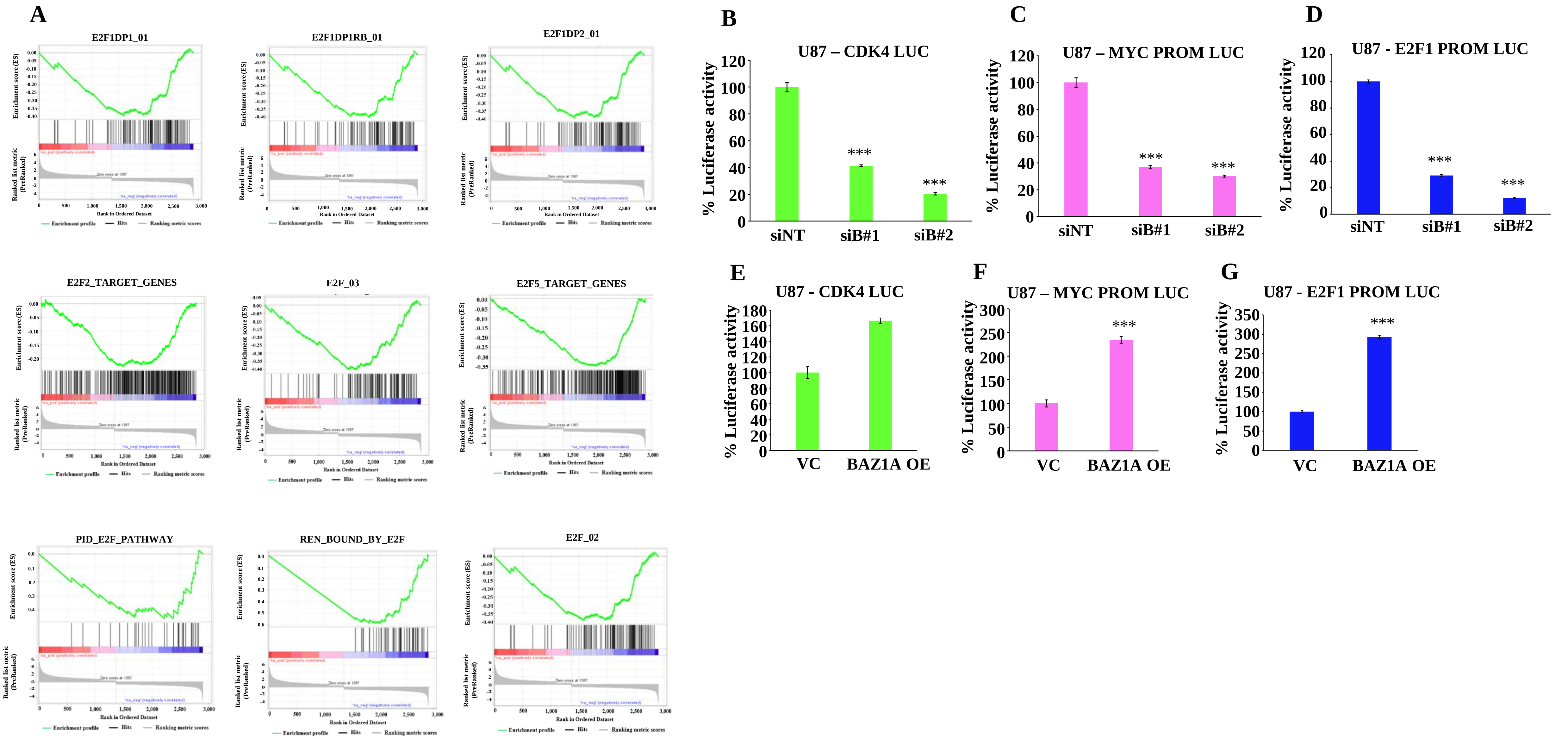
A
E2F1DP1_01
E2F1DP1RB_01
E2F1DP2_01
E2F2_TARGET_GENES
E2F_03
E2F5_TARGET_GENES
PID_E2F_PATHWAY
REN_BOUND_BY_E2F
E2F_02
Enrichment score (ES)
Ranked list metric (PreRanked)
Rank in Ordered Dataset
Enrichment profile
Hits
Ranking metric scores
B
U87 – CDK4 LUC
C
U87 – MYC PROM LUC
D
U87 - E2F1 PROM LUC
% Luciferase activity
siNT
siB#1
siB#2
***
E
U87 - CDK4 LUC
F
U87 – MYC PROM LUC
G
U87 - E2F1 PROM LUC
VC
BAZ1A OE

**Supplementary figure 6**: **BAZ1A regulates E2F transcriptional program.** **A**. Enrichment plots of E2F gene sets which were significantly depleted in BAZ1A-silenced LN229 cells. **B**, **C** and **D**. CDK4, MYC and E2F1 promoter luciferase activity measurement in BAZ1A-silenced U87 cells. Cells were harvested after 72 hours of transfection of siRNA and luciferase constructs. **E**, **F** and **G**. CDK4, MYC and E2F1 promoter luciferase activity measurement in BAZ1A overexpressed U87 cells. Cells were harvested after 24 hours of transfection of BAZ1A overexpression and luciferase constructs. Student's t-test was performed, where *:$p<0.05$,**:$p<0.01$,***:$p<0.001$, ns: non-significant.

Supplementary Figure 7; Prajapati et al., 2024

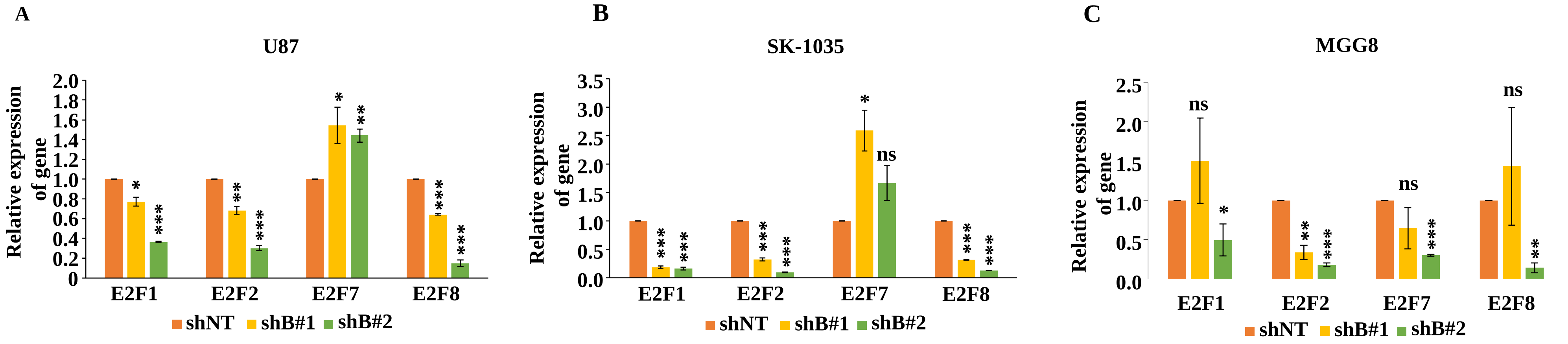

**Supplementary figure 7**: **BAZ1A regulates E2F transcriptional program. A**. RT-qPCR analysis of E2F1, E2F2, E2F7, and E2F8 transcripts in BAZ1A depleted U87 cells. **B** and **C**. RT-qPCR analysis of E2F1, E2F2, E2F7, and E2F8 transcripts in BAZ1A depleted SK-1035 and MGG8 cells. Student's t-test was performed, where *:$p<0.05$,**:$p<0.01$,***:$p<0.001$, ns: non-significant.

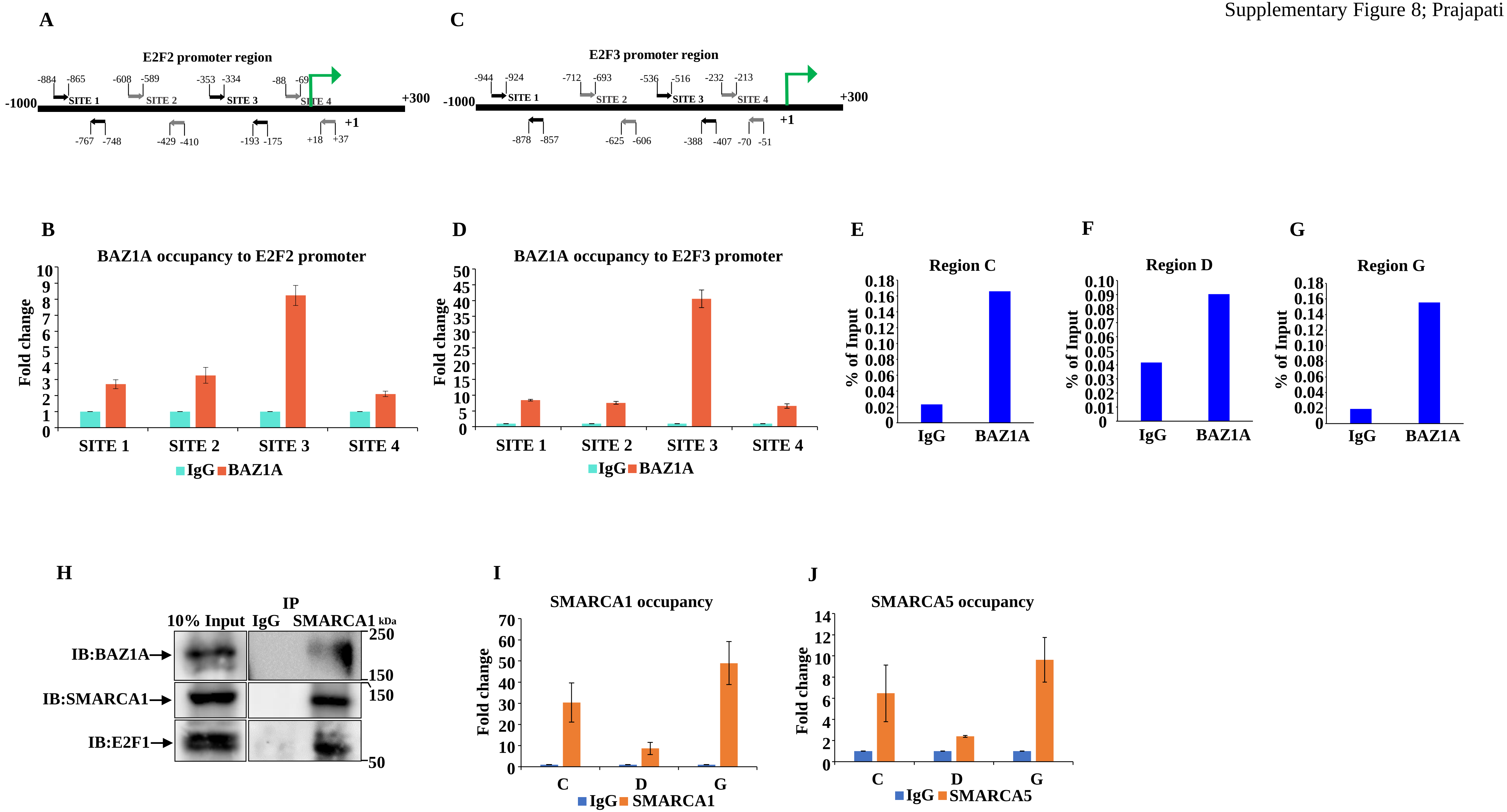
A
E2F2 promoter region
-884 -865 SITE 1
-608 -589 SITE 2
-353 -334 SITE 3
-88 -69 SITE 4
-1000
+300
+1
-767 -748
-429 -410
-193 -175
+18 +37
C
E2F3 promoter region
-944 -924 SITE 1
-712 -693 SITE 2
-536 -516 SITE 3
-232 -213 SITE 4
-1000
+300
+1
-878 -857
-625 -606
-388 -407
-70 -51
B
BAZ1A occupancy to E2F2 promoter
Fold change
SITE 1
SITE 2
SITE 3
SITE 4
IgG BAZ1A
D
BAZ1A occupancy to E2F3 promoter
Fold change
SITE 1
SITE 2
SITE 3
SITE 4
IgG BAZ1A
E
Region C
% of Input
IgG
BAZ1A
F
Region D
% of Input
IgG
BAZ1A
G
Region G
% of Input
IgG
BAZ1A
H
IP
10% Input
IgG
SMARCA1
kDa
IB:BAZ1A
IB:SMARCA1
IB:E2F1
250
150
150
50
I
SMARCA1 occupancy
Fold change
C
D
G
IgG SMARCA1
J
SMARCA5 occupancy
Fold change
C
D
G
IgG SMARCA5

**Supplementary figure 8**: **E2F1 gene expression is regulated by BAZ1A. A and C**. Schematic of E2F2 and E2F3 promoter region. **B** and **D**. ChIP assay with BAZ1A antibody followed by qPCR of the E2F2 and E2F3 promoter regions. **E**, **F,** and **G**. First ChIP with BAZ1A antibody for regions C, D and G as a positive control. **H**. HEK293T cells were transfected with BAZ1A-eGFP and HA-E2F1 and harvested after 24 hours for whole cell lysate preparation. Immunoprecipitation for SMARAC1 was performed with the lysates with specific antibodies, which were then resolved by SDS-PAGE, and immunoblotting was performed with indicated antibodies. 10% of inputs are indicated. **I** and **J**. ChIP followed by qPCR with SMARCA1 and SMARCA5 antibodies to E2F1 promoter region C, D and G. IgG was used as a control.
