## Supplementary methods for "BAZ1A, an Imitation Switch (ISWI) protein, interacts and facilitates the recruitment of E2F1 to activate the E2F transcription program"

**Supplementary information – Prajapati et al., 2024**

**Differential expression analysis**

The GlioVis database was used to obtain gene expression data from multiple GBM datasets, namely – TCGA-Agilent, TCGA-Affymetrix, TCGA-RNA-seq (RRID: SCR_003193), REMBRANDT (RRID: SCR_004704), Murat, Gravendeel, Kamoun, Grzmil, Gill and CGGA (RRID: SCR_01882). For differential expression analysis, the average of the $log_2$ transformed normalized read counts of control samples was subtracted from that of the GBM samples. Statistical significance was calculated using the Student's t-test. Genes were considered differentially expressed at $|log2FC| \geq 0.58$, and Benjamini-Hochberg adjusted p-value < 0.05.

**Correlation analysis**

To analyze the transcript correlation of BAZ1A with E2F genes and their target genes, we utilized multiple GBM datasets (including TCGA-Agilent, TCGA-Affymetrix, TCGA-RNA-seq, REMBRANDT, Murat, Gravendeel, Kamoun, Grzmil, Gill, CGGA, Oh, Philips, Freije, LeeY, Bao, Reifenberger, Donson, Ducray, Joo, Kwom, Li, Nutt, Vital, and Walsh) as well as in-house GBM samples obtained from NIMHANS. A Pearson correlation coefficient of ≥ 0.3 was used as the cutoff to classify gene pairs as positively correlated.

**RNA-sequencing**

Total RNA was extracted from shNT and shBAZ1A cells using the Trizol method. Quality of RNA-seq libraries were checked using high sensitivity D1000 ScreenTape in Agilent 2200 Tape Station system and final library quantification was done using real time PCR (quant studio & flex). Sequencing was performed with 2 * 100bp paired end reads using Novaseq 6000 (Illumina). After data generation, the integrity of the raw reads was checked and quality control was performed with FastQC ((RRID: SCR_014583 http://www.bioinformatics.babraham.ac.uk/projects/fastqc/). Human transcriptome FASTA file (GRCh38) was downloaded from GENCODE (RRID: SCR_014966) and was used to create a transcriptome index using Kallisto **(**[14]**).** Transcript-level abundance estimates of genes produced by Kallisto were converted to gene-level estimates using tximport R package (RRID: SCR_016752) **(**[15]**)**. Differential expression of genes between shNT and shBAZ1A was assessed using the negative binomial model of DESeq2 (RRID:SCR_000154) **(**[16]**).** Genes were considered differentially expressed at │log2FC│ ≥ 0.58, and Benjamini-Hochberg adjusted p-value < 0.05. The ComplexHeatmap R package was used to create heatmaps using the TPM-normalized read counts of the differentially expressed genes **(**[17]**).** The Integrative Genomics Viewer (IGV) was used to visualize the coverage of reads in both replicates of shNT and shBAZ1A using the corresponding BAM files. Volcano plot was created using ggplot2 R package ((RRID: SCR_014601) to display the differential regulation of genes ([18]).

To visualize and compare read coverage of BAZ1A, E2F1, E2F2, E2F7, and E2F8 between shNT and shBAZ1A, we utilized Broad Institute's Integrative Genomics Viewer (IGV). The BAM files generated by Kallisto were aligned to the human reference genome (GRCh38) to produce the genome tracks.

**Data availability**

The RNA seq data is submitted to Gene Expression Omnibus (GEO) (RRID: SCR_005012). As soon as the number is obtained, it will be shared.

**Gene set enrichment analysis**

Patients in the TCGA-Agilent cohort were divided into two groups based on high or low expression of BAZ1A. Differentially expressed genes were ranked according to their regulation, and the pre-ranked list was used for gene set enrichment analysis using GSEA (RRID: SCR_003199; http://www.broad.mit.edu/gsea/) with the Hallmark gene set from the Molecular Signature Database (MSigDB). A similar analysis was conducted comparing differentially expressed genes between shNT and shBAZ1A.

**Plasmids and siRNAs**

shRNA plasmids were obtained from TRC library (Sigma, IISc) – shBAZ1A#1 (TRCN0000229785), shBAZ1A#2 (TRCN0000218199), shCTNNB1#1 (TRCN0000314921), shCTNNB1#2 (TRCN0000314991), shLEF1#1 (TRCN0000020162) and shLEF1#2 (TRCN0000020163). The helper plasmids – psPAX2 (Addgene #12260; RRID:Addgene_12260), pMD2.G (Addgene #12259; RRID:Addgene_12259), and shNT (non-targeting) for lentivirus preparation is a kind gift from Dr. Subba Rao. BAZ1A overexpression construct BAZ1A-eGFP was generously provided by Dr. Akira Yasui. pCMVHA-E2F1 (Addgene #24225; RRID: Addgene_24225) was obtained from Dr. Shweta Tyagi. E2F1 PROM LUC (pGL2-AN, Addgene #20950; RRID: Addgene_20950), MYC PROM LUC (HBM-luc, Addgene #35155; RRID: Addgene_35155 ) and CDK4 PROM LUC (pBV-Luc wt MBS 1-4, Addgene #16564; RRID:Addgene_16564 ) was from Addgene. siRNA against BAZ1A used was bought from GeneX – siBAZ1A#1 (GUCUGCUAUUGUUAAAGCA) and siBAZ1A#2 (AACACUGUGAACCACAAGAUG).

**Lentivirus preparation and infection**

shRNA plasmid for specific genes and helper plasmids psPAX2 and pMD2.G were transfected together to HEK293T cells using Opti-MEM medium (Invitrogen) and lipofectamine 2000 (Invitrogen). The transfection media was replaced by fresh DMEM media containing 10% FBS after 5 hours of transfection, followed by the supernatant collection after 60 hours of transfection. The supernatant was centrifuged for 10 min at 5000 rpm, and aliquots were stored at -80°C. 0.8 and 1 million cells were seeded in 60mm culture dishes. After 16 to 18 hours, the cells were infected with aliquoted lentivirus (encoding either non-targeting shRNA or shRNA against a specific gene) along with hexadimethrine bromide (10 µg/ml Polybrene, Sigma-Aldrich, U.S.A). Post 24 hours of infection, the cells were washed with 1X PBS and replenished with fresh DMEM. Puromycin was then added at a 1 µg/ml concentration to select virus-infected cells over 48 hours. After selection, the cells were cultured in fresh DMEM and subsequently seeded for further experiments.

**Sphere formation assay**

GSCs were chemically dissociated using the Neurocult TM Chemical Dissociation Kit (Catalog #05707, STEMCELL Technologies), counted with a hemocytometer, and 50,000-60,000 cells were plated in duplicate in 6-well ultra low-attachment plates (Corning). Six hours after seeding, the cells were infected with lentivirus. 48 hours post-infection, the spheres were chemically dissociated and counted, and 5,000 cells per well were plated in duplicate in 24-well ultra-low attachment plates. After seven days, images of the spheres were captured using an Olympus microscope. Sphere diameters were measured and quantified with a specific cutoff using ImageJ software(RRID: SCR_003070).

**Limiting dilution assay**

GSCs transduced with shNT and shBAZ1A were counted and plated in 96-well low-attachment plates. For each condition, 12, 25, 50, 100, and 200 single cells were plated per well, and sphere formation was monitored for 7 to 8 days. The Extreme Limiting Dilution Assay (ELDA) software is used to plot the number of wells that do not form spheres against the number of cells plated per well (https://www.elda.at/cdscontent/?contentid=10007.854970&portal=eldaportal).

**RNA isolation, cDNA synthesis, and RT-qPCR**

Total RNA was isolated from the cells using TRI reagent (Sigma) following the manufacturer's instructions. RNA integrity was evaluated by running it on a 2% MOPS formaldehyde gel. After quantification with a Nanodrop (Thermo), 2 µg of RNA was used for cDNA synthesis with a cDNA conversion kit (Applied Biosystems). Gene knockdown or silencing was assessed by RT-qPCR using DyNamo master mix (Applied Biosystems) on a QuantStudio5 real-time PCR machine (Applied Biosystems). 20 ng of cDNA was used with gene-specific primer pairs, and ATP5G, RPL35, ACTB, and GAPDH served as internal controls. The ΔΔCT method was employed to analyze gene expression. The PCR conditions were as follows: 95°C for 5 minutes; 40 cycles of 95°C for 30 seconds, 60°C for 30 seconds, and 72°C for 30 seconds.

**Proliferation assay using MTT**

Cells infected with shNT or shRNA targeting specific genes were seeded at a density of 500 or 1000 cells per well in a 96-well plate. To quantify the actively proliferating cell population, 10 µl of MTT (0.5 mg/ml in PBS, Sigma, U.S.A.) was added to each well. Three hours after the MTT addition, the resulting formazan crystals were dissolved in 200 µl of DMSO, thoroughly mixed, and the absorbance was measured at 570 nm using an ELISA plate reader (Tecan). Statistical significance was determined using the Student's t-test.

**Viability assay**

Cells infected with shNT or shRNA targeting specific genes were seeded at a density of 20000 cells per well in a 12-well plate. Cells were harvested by trypsinization on days 1, 3, 5, and 7, and the number of viable cells was measured using a Vi-CELL XR cell viability analyzer (Beckman Coulter). The viable cells/ml of cell suspension were counted and plotted. Statistical significance was determined using the Student's t-test.

**Chemosensitivity assay**

Cells transduced with shNT or shBAZ1A were harvested after puromycin selection and seeded at a density of 500 cells per well in 96-well plates. Various concentrations of Temozolomide (TMZ) dissolved in DMSO and control (DMSO only) were used. After 72 hours of TMZ treatment, 20 µl of MTT (0.5 mg/ml) was added and incubated for three hours. The resulting formazan crystals were dissolved in 200 µl of DMSO, mixed thoroughly, and absorbance was measured at 570 nm using a Tecan reader. Absorbance was normalized to control samples. Statistical significance was determined using the Student's t-test. Similarly, shNT or shBAZ1A transduced LN229 or U87 cells were plated in 35mm dishes, and different concentrations of

TMZ were added 24 hours post-seeding. Cells were incubated with TMZ for 48 to 72 hours and harvested for apoptosis and cell cycle assays.

**Colony formation assay**

Cells treated with shNT or shRNA targeting a specific gene were seeded in six-well plates at 1500 cells/well density. The culture media were replaced every other day. After 12 days of culture, the colonies formed were fixed with 100% chilled methanol for one hour at - 20°C, followed by staining with 0.5% crystal violet for 30 minutes. Images of the colonies were captured using a Bio-Rad GelDoc, and the colonies were counted. Statistical significance was determined using the Student's t-test.

**Migration assays**

Cells were seeded in duplicate at a density of 0.2 million cells/well in 12-well plates. After 24 hours of seeding, a scratch was made with the tip end, and the media was replaced with fresh incomplete DMEM (without FBS). Images were captured at 0, 24, 48, and 72 hours after the scratch was made. Statistical significance was determined using the Student's t-test.

**Cell cycle analysis by flow cytometry**

For other cell cycle experiments, cells were harvested at specific time points. Following two washes with 1X PBS, cells were fixed with chilled ethanol (70%) and incubated overnight at - 20°C. Subsequently, the fixed cells were washed twice with 1X PBS, treated with RNAse A (20 µg/ml) for 2 hours at 37°C, and stained with Propidium Iodide (PI) at 10 µg/ml concentration. The cells were then analyzed using flow cytometry with a FACS VERSE instrument (BD Biosciences) and analyzed using FACSsuite software (BD Biosciences).

**BrdU incorporation assay using flow cytometry**

For the BrdU incorporation assay, cells were incubated with 10 µM BrdU for 20 minutes at 37°C in the incubator. After washing with 1X PBS three times, cells were fixed in 70% chilled ethanol and left overnight at -20°C. Following a PBS wash, DNA was denatured using 0.5% Triton X-100 containing hydrochloric acid (2N) for 30 minutes at room temperature. Once neutralized with PBS, cells were blocked with 0.5% Triton X-100 containing 0.5% BSA for 30 minutes. Subsequently, cells were incubated with anti-BrdU antibody (BrdU (Bu20A), sc-20045, RRID:AB_626767 1:100) for 2 hours at room temperature, followed by conjugation with a secondary antibody (Invitrogen, Alexa 488 anti-mouse IgG, catalog #A11029) for one hour at RT. After careful washing, cells were treated with RNase A (100 ng/ml) for 30 minutes to one hour. Propidium iodide (50 μg/ml) was added to the cells, and they were analyzed using flow cytometry with a FACSVERSE instrument (BD Biosciences) and FACSsuite software (BD Biosciences).

**Annexin V-FITC (Fluorescein isothiocyanate) and PI staining**

Apoptosis was evaluated using Annexin V-FITC/PI double staining. According to the manufacturer's instructions, flow cytometry-based analysis was conducted using the Annexin V-FITC Apoptosis kit from MACS Miltenyi Biotech. The early and late apoptotic cell percentages were determined and plotted as the total apoptotic cells per condition. Statistical significance was determined using the Student's t-test.

**Luciferase assay**

After 16-18 hours of seeding, cells were co-transfected with Luciferase reporter constructs along with either vector control or overexpression constructs or with siRNAs. Following 24 or 72 hours of transfection, cells were harvested and lysed in reporter lysis buffer (Promega, #E3971). An equal amount of protein was utilized for the luciferase assay, using luciferase assay reagent (Promega, #E1483) and a luminometer (Berthold detection system, Sirius). Statistical significance was determined using the Student's t-test.

**Immunoblotting**

Cell lysates were prepared using RIPA lysis buffer (containing 50 mM Tris HCl, 150 mM NaCl, 1.0% NP-40, 0.5% Sodium Deoxycholate, 1.0 mM EDTA, 0.1% SDS, and 0.01% sodium azide at pH 7.4), supplemented with Sodium orthovanadate (1 mM), Sodium Fluoride (1 mM), Phenylmethylsulfonyl fluoride (1 mM), and Protease inhibitor cocktail (Sigma, #S8830). Cells were lysed by scratching the cells dissolved in RIPA buffer, followed by incubation on ice for 30 minutes and centrifugation at 14,000 rpm at 4°C for 30 minutes. Protein quantification was performed using Bradford reagent in an ELISA plate reader (Tecan). Protein lysates were denatured for 15 minutes at 95°C. Subsequently, 10% SDS-PAGE was run, and the resolved proteins were transferred to a 0.45-micron PVDF membrane using the semi-dry transfer method (Biorad). Non-specific proteins were blocked by incubating the membrane with 5% non-fat dried milk in TBST buffer (25 mM Tris-HCl pH 7.5, 150 mM NaCl, 0.05% Tween 20) for 1 hour at RT. After washing the membrane with TBST buffer once, it was probed with specific primary antibody overnight at 4°C. Following three washes with TBST buffer, the membrane was incubated with HRP-conjugated secondary antibody (Invitrogen, Goat anti-rabbit IgG - catalog#31460 (RRID: AB_228341) and Goat anti-mouse IgG - catalog#31430

(RRID: AB_228307)) at RT for 1-2 hours. After three washes with TBST buffer, chemiluminescent signals were detected using Clarity ECL Western Blotting Substrate (BioRad) in a chemiluminescence imager (Chemidoc Touch, Biorad).

**Immunoprecipitation assays**

Immunoprecipitation experiments were conducted following standard protocols. Cells were scraped from the culture dish in Pierce IP lysis buffer (containing 25 mM Tris-HCl pH 7.4, 150 mM NaCl, 1 mM EDTA, 1% NP-40, and 5% glycerol), supplemented with Sodium orthovanadate (1 mM), Sodium Fluoride (1 mM), Phenylmethylsulfonyl fluoride (1 mM), and Protease inhibitor cocktail (Sigma, #S8830), and lysed at 4°C for 30 minutes with agitation. Debris was removed by centrifugation of the lysate at 14,000 rpm at 4°C. 1 mg of protein lysate was incubated with 5 μg of antibody or control IgG antibody for 18-20 hours at 4°C. 30 ul of protein G-conjugated magnetic beads (Invitrogen, catalog #10004D) were then added to each set and incubated for 4 hours at 4°C to capture the immune complexes. Beads were washed three times with ice-cold Pierce IP lysis buffer by gentle rotation at 4°C for 5 minutes each. The immunoprecipitated proteins were denatured at 95°C for 15 minutes in 6X Laemmli buffer and resolved by 10% SDS-PAGE, followed by transfer to a PVDF membrane (MERCK, Cat #IPVH00010). The membrane was blocked with TBST buffer containing 5% non-fat dried milk for 1 hour at room temperature. After three washes with TBST buffer, the membrane was probed with primary antibody overnight at 4°C. Following primary antibody incubation, the membrane was probed with HRP-conjugated secondary antibodies (Invitrogen, Goat anti-rabbit IgG - catalog#31460 and Goat anti-mouse IgG - catalog#31430), or light-chain specific HRP-conjugated secondary antibodies (Anti-mouse IgG, light-chain specific, #D3V2A, rabbit mAb (RRID:AB_2799549) and Mouse anti-rabbit IgG, light-chain specific, #D4W3E, mouse((RRID: AB_2799281)) at room temperature for 1-2 hours. Chemiluminescent signals were detected using Clarity ECL Western Blotting Substrate (BioRad, catalog #170-5061) in a chemiluminescence imager (Chemidoc Touch, Biorad).

**Chromatin Immunoprecipitation (ChIP)**

Chromatin isolation from LN229 cells was performed following the manufacturer's protocol (CST; Cat no. 9003). Briefly, cells were cross-linked with 1% formaldehyde and incubated at RT for 10 minutes. The reaction was then quenched by adding 125 mM glycine and incubating for 5 minutes at room temperature. Nuclei were subsequently prepared using specific buffers and digested with MNase, followed by sonication to generate chromatin fragments ranging from 150 to 900 bp. The sheared chromatin was then incubated with 2 µg of the specific antibody/antibodies for 16 hours, followed by incubation with 30ul of Protein G magnetic beads (Invitrogen) for 4 hours at 4°C. An equal amount of IgG antibody was used as a negative control. Chromatin DNA was eluted using the phenol-chloroform isolation method, and the eluted DNA was utilized for quantitative PCR with promoter-specific primers to amplify the desired region of the promoter. The PCR conditions were as follows: 95°C for 5 minutes; 40 cycles of 95°C for 30 seconds, 60°C for 30 seconds, and 72°C for 30 seconds. Fold enrichment over IgG was calculated.

**ChIP ReChIP or sequential Chromatin Immunoprecipitation**

Chromatin was prepared according to the manufacturer's instructions (CST; Cat no. 9003) as described above. From the prepared chromatin, 5% input was set aside separately. The remaining chromatin was divided into two portions for individual ChIP experiments with BAZ1A and E2F1 antibodies. Each ChIP experiment was conducted with 20 million LN229 cells. Following the final wash of the beads with wash buffers, the bound chromatin was eluted with 75 µl of TE buffer (10 mM Tris-HCl pH 8, 2 mM EDTA) containing 10 mM DTT by incubating the beads at 37°C for 1 hour on a thermal shaker (to denature the first antibody used). The eluted supernatant was collected by centrifugation at 800g for 2 minutes. This eluted chromatin supernatant was diluted 70-fold with ChIP buffer to prevent interference from DTT on the second antibody. Each diluted eluate was divided into two portions for ReChIP experiments—one for the second specific antibody and another for the IgG control. The final chromatin was eluted (using CST; Cat no. 9003), and DNA purification was performed using the standard phenol-chloroform method. The eluted DNA was subjected to qPCR using E2F1 promoter-specific primers to amplify the desired region of the promoter. The PCR conditions were as follows: 95°C for 5 minutes; 40 cycles of 95°C for 30 seconds, 60°C for 30 seconds, and 72°C for 30 seconds. Percent input was calculated based on the DNA from the 5% input sample.

**DNaseI hypersensitivity assay**

shNT or shBAZ1A transduced LN229 cells were cross-linked using 1% formaldehyde and subsequently quenched with 125 mM glycine. After washing three times with chilled 1X PBS, the cells were scraped in 1X PBS containing protease inhibitor supplements and centrifuged at 2500g for 10 minutes at 4°C. The collected cells were then incubated with nuclei lysis buffer (containing 50 mM Tris HCl pH 7.4, 1% SDS, 10 mM EDTA pH 8; supplemented with protease inhibitors) for 10 minutes on ice to lyse the nuclei and sonicated to generate chromatin fragments. Following a 5-minute incubation on ice, the chromatin supernatant was collected by centrifugation at 12000 rpm for 15 minutes at 4°C. 2 ug of chromatin were measured by nanodrop from each condition and treated with 0, 0.5, 1, and 2 units of DNaseI enzyme (Promega #M6101) for 3 minutes at 37°C. The reaction was stopped by incubating the reaction mixture with stop buffer (20 mM EGTA pH 8.0, promega #M6101) and keeping on ice for 5 minutes. After treating the reaction mixture with RNase A (10 mg/ml) for 2 hours at 37°C, proteinase K (10 mg/ml) was added and incubated at 55°C for 2 hours. DNA purification was carried out using the standard phenol-chloroform method, and an equal volume of DNA was used for qPCR to amplify the desired promoter region.

**Orthotopic xenograft mouse model**

The Institutional Animal Ethics Committee (IAEC) approved all animal procedures (CAF/Ethics/925/2022). U-87 MG-Luc2 cells were transduced with shNT and shB#1 lentivirus. After 72 hours post-infection, cells were trypsinized, and an equal number of cells were taken for orthotopic injection. Specifically, 0.3 million cells were stereotaxically injected into the hippocampus of 5-6-week-old athymic nude mice (RRID: IMSR_JAX:002019) using a stereotaxic apparatus (Kopf and RWD) with the coordinates: AP = +2.0, ML = +1.5, DV = 2.5. The animals were subjected to bioluminescence imaging at regular intervals, where the total photon flux (photon/s) was measured and plotted. Survival of the shNT (n=16) and shBAZ1A (n=16) groups was monitored and plotted using GraphPad Prism 8 (RRID: SCR_002798). The difference in survival between the two groups was assessed using the Mantel-Cox log-rank test. (AP=Anterior-Posterior, ML=Medial-Lateral, DV=Dorsal-Ventral.)

***In-vivo* imaging**

For bioluminescence imaging, D-Luciferin Sodium Salt (GOLDBIO, CAS#103404-75-7, LUCNA-1G) was intraperitoneally injected into each mouse at a dosage of 150 mg/kg body weight. Following injection, mice were anesthetized using gas anesthesia with isoflurane and then positioned in the IVIS machine for imaging. *In vivo* bioluminescence imaging was conducted using the PerkinElmer IVIS Spectrum system.

**Cryo-sectioning of fixed mouse brain**

Transcardiac perfusion was conducted on selected mice using 1X PBS followed by a 4% paraformaldehyde solution. Subsequently, the mouse brains were harvested and immersed in 4% PFA for 24 hours, followed by storage in a 30% sucrose solution for 48 hours. Poly-freeze solution (Sigma, #35059990) was utilized to embed the mouse brains, and the embedded brains were then sectioned into 30 µm thick slices using an RWD Cryostat. These sections were preserved at -80°C in Tissue Cutting Solution (composed of 0.1 M phosphate buffer, ethylene glycol, and glycerol).

**Immunohistochemistry (IHC)**

Brain sections were removed from Tissue Cutting Solution for immunofluorescence analysis, placed in 12-well plates, and thoroughly washed with 1X PBST. Subsequently, the tissues were incubated in a permeabilization solution (1X PBS and 0.25% Triton-X 100) for one hour at RT followed by blocking using a solution containing 1X PBS, 1% BSA, 0.3% Triton-X 100, and 10% goat serum for 2-3 hours. After two washes with PBST, primary antibodies (Ki-67, Abcam, catalog #ab16667, RRID:AB_302459; 1:100, and E2F1, ThermoFisher Scientific, catalog #32-1400, RRID:AB_86934; 1:100) prepared in blocking solution were added to the tissues in a 96-well plate and incubated overnight at 4°C. Following overnight incubation, the tissues were washed twice with PBST and then incubated with secondary antibodies (Invitrogen, Alexa 488 anti-mouse IgG, catalog #A11029 (RRID: AB_2534088), and Alexa 594 anti-rabbit IgG, catalog #A11037 (RRID: AB_2534095)) prepared in blocking solution at RT for 2 hours. Subsequently, the tissues were stained with DAPI (100 ng/ml) for 30 minutes and washed twice with PBST. Finally, the tissues were individually mounted on glass slides using ProLong™ Glass Antifade Mountant (Invitrogen, #P36980) and covered with coverslips. Images were acquired using Leica Falcon (10X magnification for whole brain images) and Andor Dragonfly (40X magnification for tissue sections).

**Hematoxylin and Eosin staining (H and E)**

Cryosections of mouse brain tissues were transferred onto glass slides coated with poly-L-lysine and incubated at 42°C for 48 hours. The slides were immersed in 70% ethanol for one minute, followed by staining with hematoxylin for 4 minutes. Subsequently, they were rinsed with distilled water for one minute and stained with eosin for one minute. After another rinse with distilled water, the tissues were differentiated by immersion in ethanol. They were then rewashed with water before being mounted using an xylene-based DPX mounting medium and covered with coverslips. Images were captured using a Lawrence and Mayo digital microscope (0.7X magnification) and a Nikon microscope (20X magnification).
